# Glyco-phenotyping of mutants of *Lacticaseibacillus paracasei* by lectin microarray

**DOI:** 10.1101/2024.09.11.612535

**Authors:** Emi Suzuki, Masaki Serata, Tomoyuki Sako, Sumie Sato, Tohru Iino, Hiroaki Tateno, Jun Hirabayashi

**Affiliations:** Yakult Central Institute, Yakult Honsha Co., Ltd., 5-11 Izumi, Kunitachi-shi, Tokyo 186-8650, Japan; Cellular and Molecular Biotechnology Research Institute, National Institute of Advanced Industrial Science and Technology (AIST), Tsukuba Central 6, 1-1-1 Higashi, Tsukuba, Ibaraki 305-8566, Japan; Tokai National Higher Education and Research System, Nagoya University, Institute of Glyco-core Research (IGCORE)

## Abstract

We previously identified a gene cluster of *Lacticaseibacillus paracasei* strain Shirota (YIT 9029) for cell surface long-chain polysaccharides (LCPS-1) biosynthesis, which modulates YIT 9029 activity to induce cytokine production in immune cells, and showed that a lectin microarray can be useful for distinguishing the profile of bacterial cell-surface polysaccharide (PS) structures. Therefore, we isolated disruptive mutant strains of 51 genes predicted to be involved in cell wall PS biosynthesis in YIT 9029. Their binding profiles to lectins in conjunction with their binding abilities to YIT 9029-specific monoclonal antibody (MAb) were compared. The mutants defective in binding to the MAb all had defects within the *cps1* gene cluster. Some mutants partially bound to MAb, indicating that these genes may influence the synthesis and maturation of LCPS-1. Advanced lectin microarray analyzed the cell surface glycosylation properties of YIT 9029 and its mutants. YIT 9029 bound to a rhamnose-specific lectin CSA, and three additional lectins including an O-glycan binder (rDiscoidin II) and two mannose binders (rOrysata and rBanana). Lectin binding specificity was confirmed by a gene complementation assay for the *cps1C* gene and a carbohydrate inhibition assay. When the binding profiles of individual *cps1A* through *cps1J* knockout mutants were compared, typical and specific binding profiles patterns were observed, in which some similarities in the functions of each gene could be predicted. In conclusion, the combined use of lectin microarray and a YIT 9029 mutant strain library is a powerful tool for identifying unknown bacterial gene functions related to cell surface glycome.

**IMPORTANCE:** Previously, only a limited number of methods have been available for studying mutations in bacterial cell surface polysaccharide structures in relation to gene function. In this study, we focused on the lectin-binding properties of *Lacticaseibacillus paracasei* YIT 9029 (wild-type; WT) and investigated the lectin-binding capabilities of 51 cell wall biosynthesis gene disruption strains using lectin microarrays. The results indicated that lectin-binding properties in gene-disrupted strains varied significantly with the presence or absence of LCPS-1, ranging from similar to WT to distinctly different. The use of lectin microarrays in conjunction with the YIT 9029 mutant library has been shown to be a highly effective method for identifying the functions of unknown bacterial genes related to cell-surface glycomes. This innovative approach to glycophenotyping allows for the determination of cell wall glycomes associated with bacterial gene functions using lectin microarrays.

## INTRODUCTION

Bacterial cell surface components, such as polysaccharide (PS) capsules, glycoproteins, or glycolipids, comprises a thick peptidoglycan (PG) layer that surrounds the cytoplasmic membrane (1) and are important signaling factors that trigger various host responses, including pathogenesis, host-microbe interaction, immune modulation, and symbiosis (2). Bacterial cell walls have unique structures related to these phenomena (1, 3). Carbohydrates in the form of capsular polysaccharides (CPS) in Gram-positive bacteria and/or lipopolysaccharides (LPS) in Gram-negative bacteria are the major components on the surface of bacteria. The cell wall of Gram-positive bacteria is a complex assembly of glycopolymers, teichoic acids (TA), and proteins. These molecules possess a thick PG layer surrounding the cytoplasmic membrane (1). Bacterial cell-surface glycans differ substantially between species and strains (4). Moreover, the cell wall PS of lactic acid bacteria (LAB) contains rhamnose (Rha) and the PS structures are highly diverse, depending on the bacterial strain (5, 6).

LAB are industrially important microorganisms for fermented food production. Many LAB strains have been reported to exert beneficial effects through immune modulation of host cells. *Lacticaseibacillus paracasei* (formerly *Lactobacillus casei*) strain Shirota (YIT 9029) exerts immunomodulatory activities in vitro (7, 8) and in humans (9–16). In addition, YIT 9029 enhances NK cell activity in healthy volunteers after regular oral feeding with intact cells (17). However, the structure and molecules of YIT 9029 responsible for these activities remain unclear.

YIT 9029 possesses two types of CPS: long-chain polysaccharides (LCPS-1 (18) and LCPS-2 (19) (formerly called PS-1 and PS-2, respectively). In previous studies, we first focused on the role of CPS in the immune modulation activities of this bacterium and revealed that certain gene-knockout strains defective in producing LCPS-1 had altered immune modulating activities toward cultured mouse macrophage-like cells (RAW264.7 and J774.1 cells) suggesting that the PS moieties of cell surface structures, including LCPS-1 (20) and LCPS-2 (19), play an important role in YIT 9029 immune modulation activities. To clarify and compare the surface structural characteristics of YIT 9029 and its mutants with different immune modulation activities, we introduced a novel approach for determining the binding profiles of these cells to lectins using liquid-phase lectin microarray technology. From this analysis, we successfully detected structural alterations on the YIT 9029 cell surface in mutants with defects in certain possible glycosylation enzymes (20).

In this study, we aimed to identify the genes of YIT 9029 that are possibly involved in the biosynthesis of genes or influence the structure of the cell surface PS moieties of the cell wall of YIT 9029 by collecting as many genes as possible and by determining the lectin-binding profiles of the gene-disrupted mutant cells in the lectin microarray system. Although databases of human and bacterial glycosyltransferase genes are being developed each year, it is still difficult to determine the role of a particular gene involved in cell wall biosynthesis. Therefore, it would be worthwhile to systematically correlate the changes in the binding profiles of these mutants with lectin probes. We identified all possible genes for the biosynthesis and modification of the cell-wall PS of YIT 9029 based on the similarity of the gene sequences with known proteins, constructed gene knockout mutants, and employed these mutants in the liquid-phase lectin microarray system.

Through this approach using lectin microarray technology, we hoped to gain new insights into the cell surface structures to evaluate other reactivity with YIT 9029-specific monoclonal antibody (MAb) (21) and the characteristics of the bacterial strain YIT 9029.

## MATERIALS AND METHODS

Criteria for the selection of genes from YIT 9029 and annotation of the genes The genes predicted to be involved in the biosynthesis of cell wall-associated PSs of YIT 9029 were selected from the in-house genome laboratories data (unpublished results; Table 1 left). These were based on the similarity of at least 20% of the amino acid sequences with other bacterial proteins known or predicted to be involved in extracellular or cell wall PSs biosynthesis in the GenBank database. In addition, genes were selected to contain motifs for glycosyl transferases and glycosylation enzymes in the genome of *L. casei* BL23 (22). The protein ID are listed in Table 1 (rightmost).

**Table 1.**
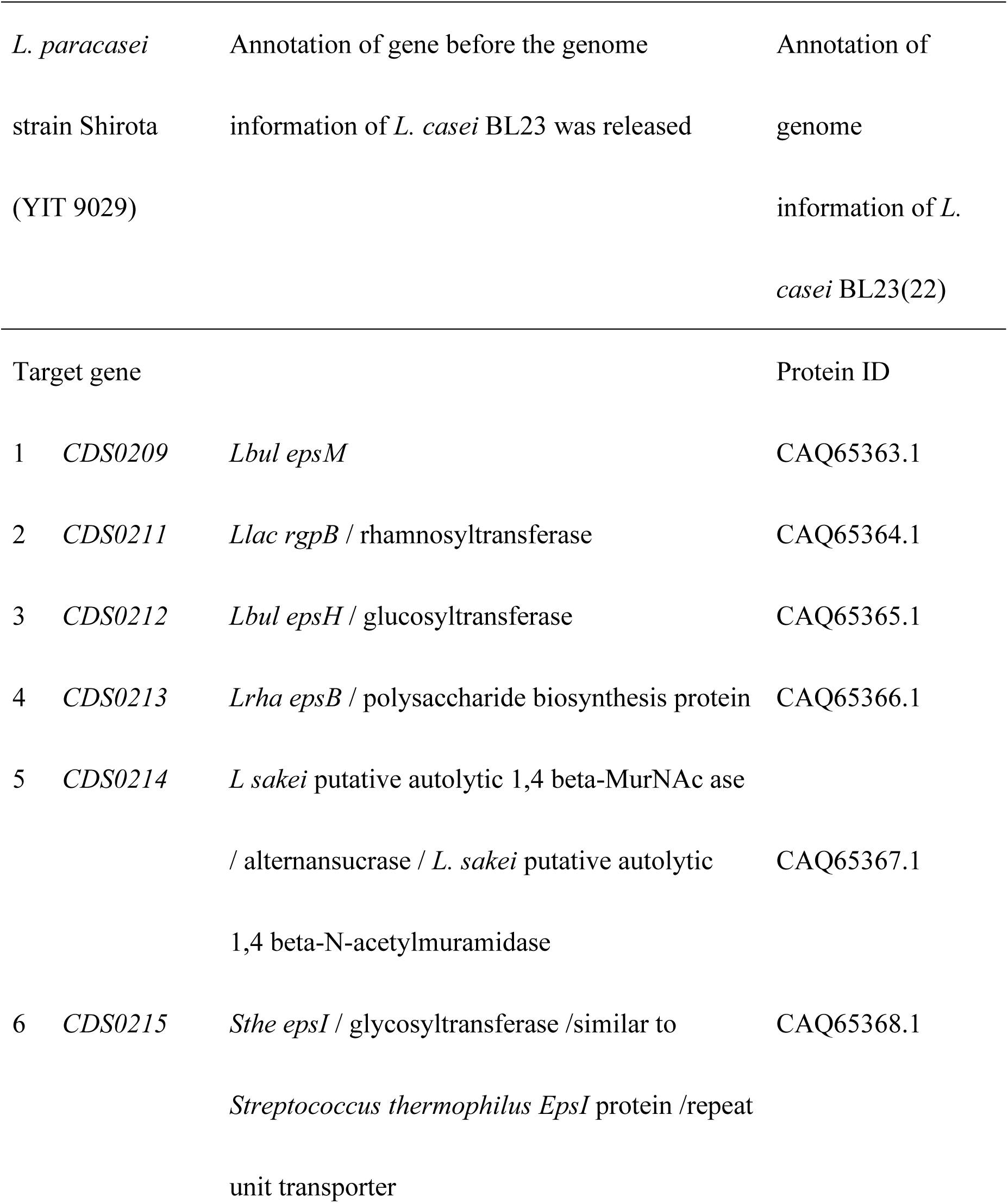
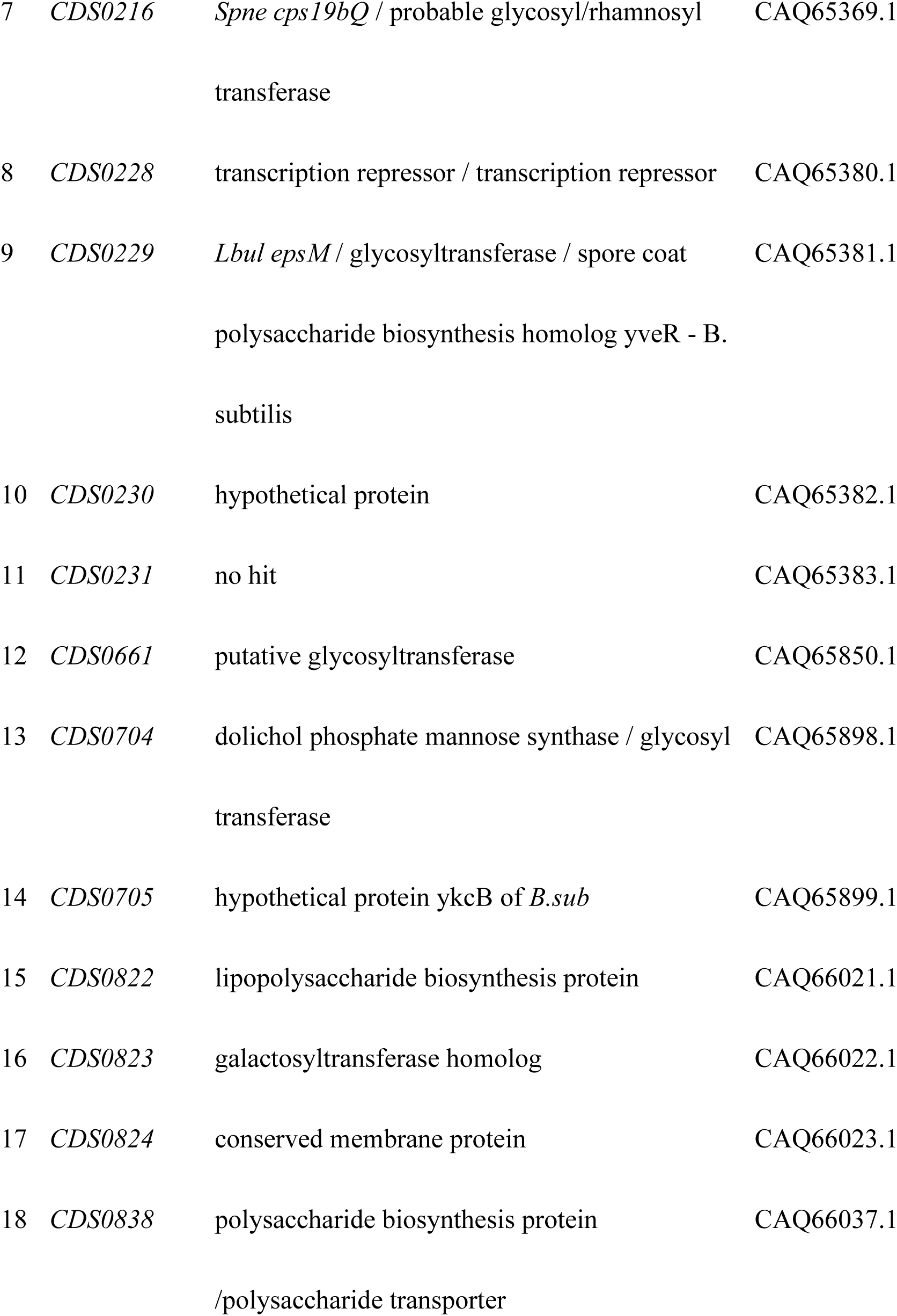
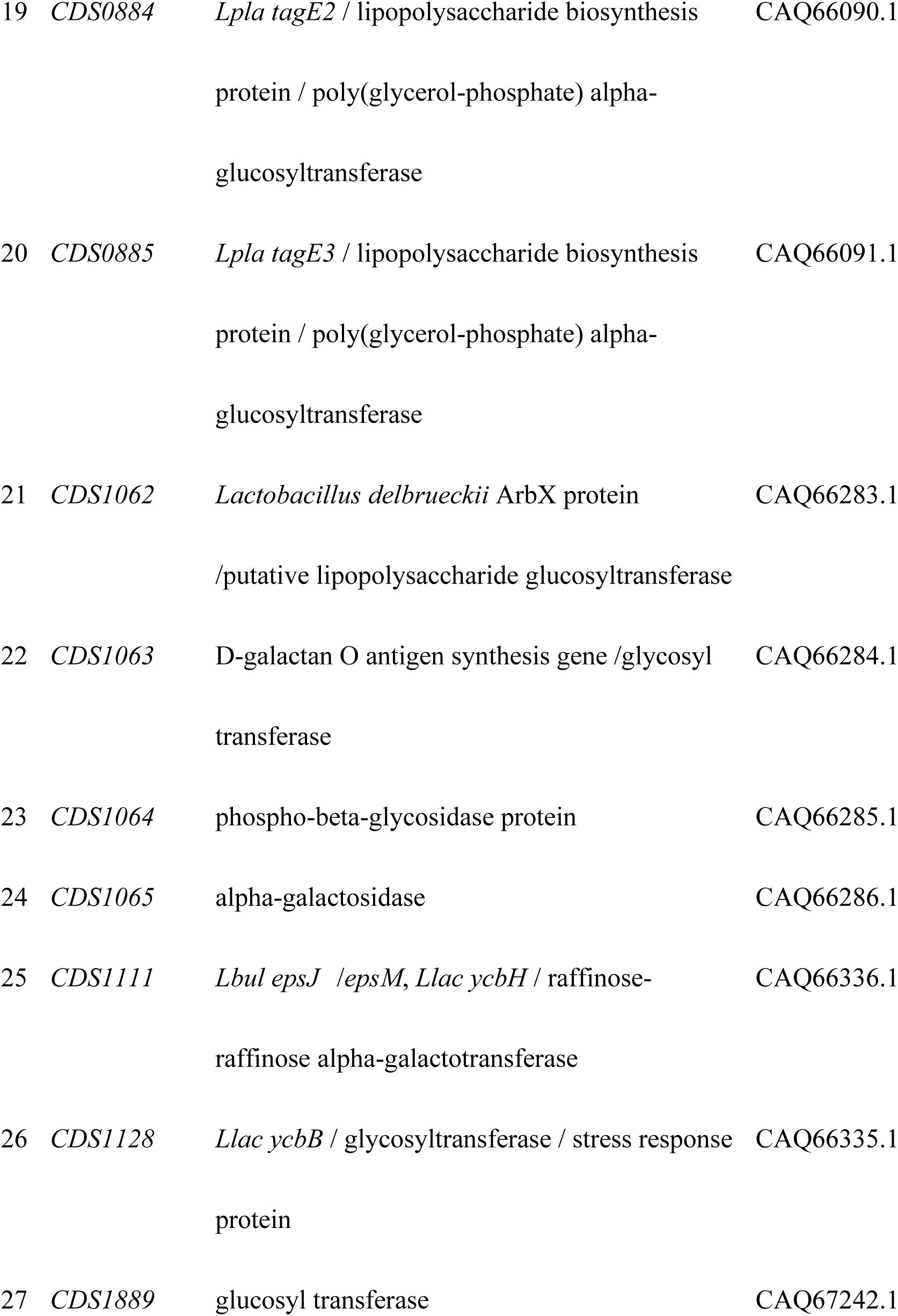
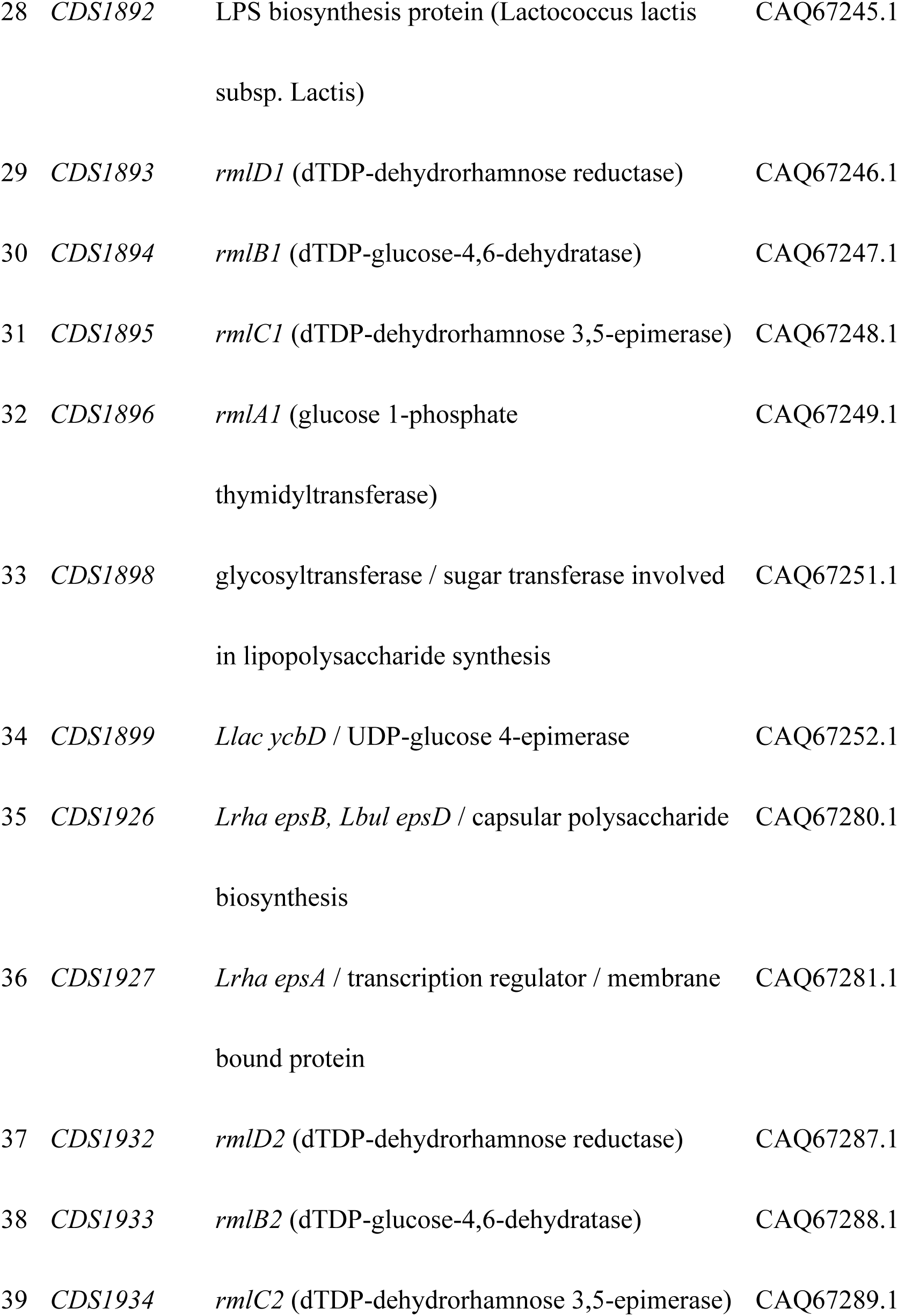
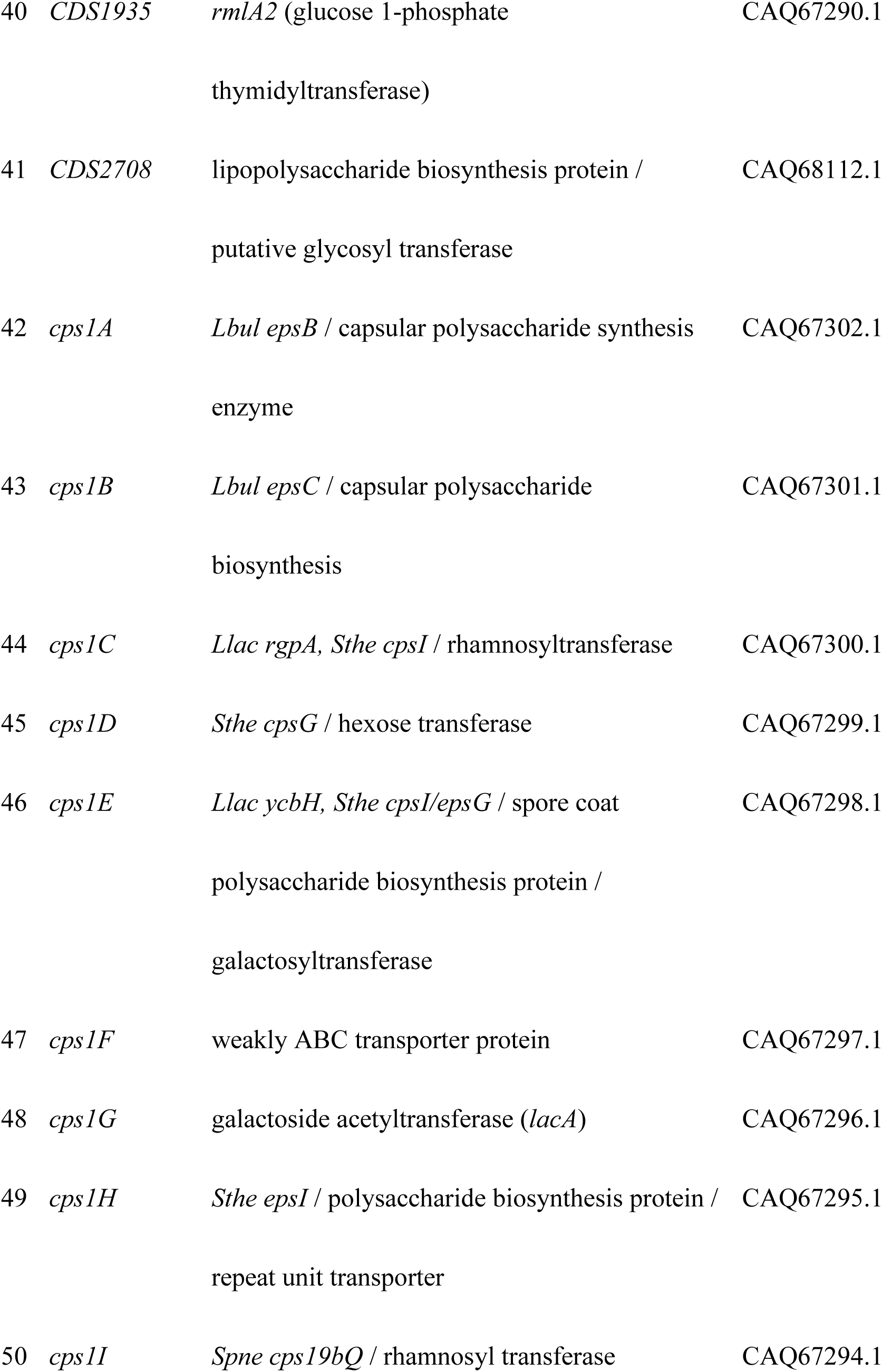
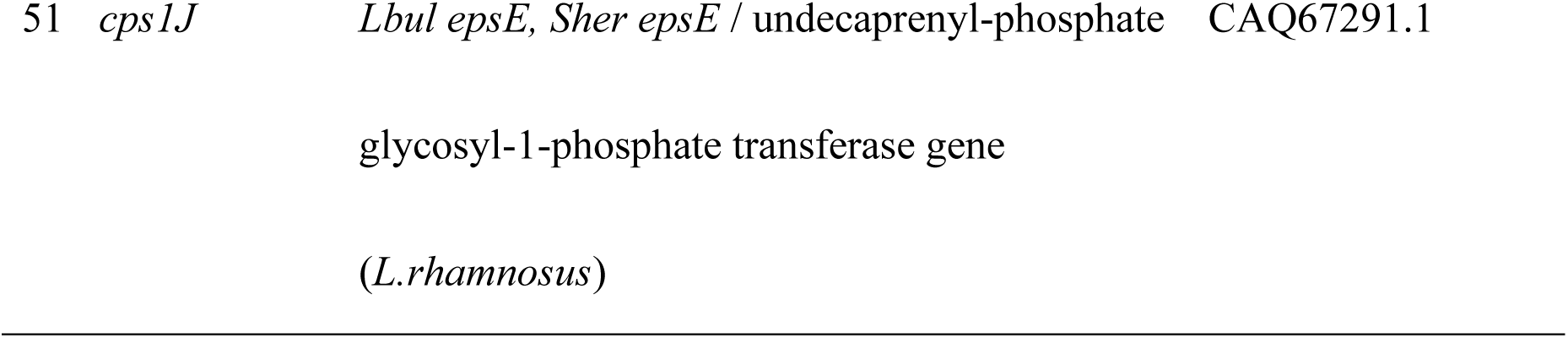
The gene library of *L. paracasei* Shirota (YIT 9029) annotated from other lactic acid bacteria and *L. casei* BL23. (**22**) **via amino acid sequence similarities of the cell wall biosynthesis related genes.**

### Bacterial strains and plasmids used in this study

The bacterial strains and plasmids used in this study are listed in Table 2. Fifty-one gene knockout mutants of YIT 9029 were employed for the analysis; some were newly constructed as described below, while others were from a previous study (20). *L. casei* ATCC 334 (YIT 0180), a neotype strain of *L. casei* (23) was purchased from the American Type Culture Collection (Manassas, VA). We tested four strains from our laboratory collection with different antibody reactivities: YIT 9021 (24), YIT 9022, YIT 9036, and YIT 9037. *Escherichia coli* JM109 was purchased from Toyobo Co. Ltd. (Osaka, Japan) as competent cells for DNA transformation.

**Table 2.** Bacterial strains used in this study.

| Strain or plasmid |  | Relevant genotype | Source or reference |
| --- | --- | --- | --- |
| Strains |  |  |  |
| *L. paracasei* |  |  |  |
| Shirota |  |  |  |
|  | YIT 9029 | Wild type | Our collection |
| 1 | $\Omega 0209$ | Insertion in *CDS0209* gene | This study |
| 2 | $\Omega 0211$ | Insertion in *CDS0211* gene | This study |
| 3 | $\Omega 0212$ | Insertion in *CDS0212* gene | This study |
| 4 | $\Omega 0213$ | Insertion in *CDS0213* gene | This study |
| 5 | $\Omega 0214$ | Insertion in *CDS0214* gene | This study |
| 6 | $\Omega 0215$ | Insertion in *CDS0215* gene | This study |
| 7 | $\Omega 0216$ | Insertion in *CDS0216* gene | This study |
| 8 | $\Omega 0228$ | Insertion in *CDS0228* gene | This study |
| 9 | $\Omega 0229$ | Insertion in *CDS0229* gene | This study |
| 10 | $\Omega 0230$ | Insertion in *CDS0230* gene | This study |
| 11 | $\Omega 0231$ | Insertion in *CDS0231* gene | This study |
| 12 | $\Omega 0661$ | Insertion in *CDS0661* gene | This study |
| 13 | $\Omega 0704$ | Insertion in *CDS0704* gene | This study |
| 14 | $\Omega 0705$ | Insertion in *CDS0705* gene | This study |
| 15 | $\Omega 0822$ | Insertion in *CDS0822* gene | This study |
| 16 | $\Omega 0823$ | Insertion in *CDS0823* gene | This study |
| 17 | $\Omega 0824$ | Insertion in *CDS0824* gene | This study |
| 18 | $\Omega 0838$ | Insertion in *CDS0838* gene | This study |
| 19 | $\Omega 0884$ | Insertion in *CDS0884* gene | This study |
| 20 | $\Omega 0885$ | Insertion in *CDS0885* gene | This study |
| 21 | $\Omega 1062$ | Insertion in *CDS1062* gene | This study |
| 22 | $\Omega 1063$ | Insertion in *CDS1063* gene | This study |
| 23 | $\Omega 1064$ | Insertion in *CDS1064* gene | This study |
| 24 | $\Omega 1065$ | Insertion in *CDS1065* gene | This study |
| 25 | $\Omega 1111$ | Insertion in *CDS1111* gene | This study |
| 26 | $\Omega 1128$ | Insertion in *CDS1128* gene | This study |
| 27 | $\Omega 1889$ | Insertion in *CDS1889* gene | This study |
| 28 | $\Omega 1892$ | Insertion in *CDS1892* gene | This study |
| 29 | $\Omega 1893$ | Insertion in *CDS1893* gene | This study |
| 30 | $\Omega 1894$ | Insertion in *CDS1894* gene | This study |
| 31 | $\Omega 1895$ | Insertion in *CDS1895* gene | This study |
| 32 | $\Omega 1896$ | Insertion in *CDS1896* gene | This study |
| 33 | $\Omega 1898$ | Insertion in *CDS1898* gene | This study |
| 34 | $\Omega 1899$ | Insertion in *CDS1899* gene | This study |
| 35 | $\Omega 1926$ | Insertion in *CDS1926* gene | This study |
| 36 | $\Omega 1927$ | Insertion in *CDS1927* gene | This study |
| 37 | $\Delta 1932$ | Deletion in *rmlD2* gene | This study |
| 38 | $\Omega 1933$ | Insertion in *rmlB2* gene | This study |
| 39 | $\Omega 1934$ | Insertion in *rmlC2* gene | This study |
| 40 | $\Omega 1935$ | Insertion in *rmlA2* gene | This study |
| 41 | $\Omega 2708$ | Insertion in *CDS2708* gene | This study |
| 42 | $\Omega cps1A$ | Insertion in *cps1A* (*CDS1945*) gene | (20) |
| 43 | $\Omega cps1B$ | Insertion in *cps1B* (*CDS1944*) gene | (20) |
| 44 | $\Delta cps1C$ | Deletion in *cps1C* (CDS1943) gene | (20, 27) |
| 45 | $\Omega cps1D$ | Insertion in *cps1D* (CDS1942) gene | (20) |
| 46 | $\Omega cps1E$ | Insertion in *cps1E* (CDS1941) gene | (20) |
| 47 | $\Omega cps1F$ | Insertion in *cps1F* (CDS1940) gene | (20) |
| 48 | $\Omega cps1G$ | Insertion in *cps1G* (CDS1939) gene | (20) |
| 49 | $\Omega cps1H$ | Insertion in *cps1H* (CDS1938) gene | (20) |
| 50 | $\Omega cps1I$ | Insertion in *cps1I* (CDS1937) gene | (20) |
| 51 | $\Omega cps1J$ | Insertion in *cps1J* (CDS1936) gene | (20) |
| 52 | $\Delta cps1A/cps1A$ | Complementary of WT-*cps1A* into<br>$\Delta cps1A$ | (20) |
| 53 | $\Delta cps1C/cps1C$ | Complementary of WT-*cps1C* into<br>$\Delta cps1C$ | (20) |
| 54 | YIT 9021 | Our laboratory collection | (24) |
| 55 | YIT 9022 | Our laboratory collection | This study |
| 56 | YIT 9036 | Our laboratory collection | This study |
| 57 | YIT 9037 | Our laboratory collection | This study |
|  | YIT 0180<sup>T</sup> | *L. casei* neotype strain | ATCC 334 |

*L. paracasei* (formerly *L. casei*) strain Shirota (YIT 9029) is a commercial strain. The YIT number shows the culture collection preserved in the Microbiological Research Department of the Yakult Central Institute of Tokyo, Japan. Mutants were produced from YIT 9029, and $\Omega$ shows a single cross over mutant. The CDS number shows the gene that encodes the YIT 9029 protein. *L. casei* ATCC 334 is the neotype strain of *L. casei* YIT 0180, which was purchased from the American Type Culture Collection (Manassas, VA). *Escherichia coli* JM109 was purchased from Toyobo Co. Ltd. (Osaka, Japan) as competent cells for DNA transformation.

### Gene manipulation of YIT 9029 based on the homologous recombination principle

Forty-nine mutants of YIT 9029 designated “Ω” were produced by insertion of a plasmid with the respective truncated gene fragment deleting both N- and C-terminal coding regions as described previously (Table 2). The synthetic primers used to amplify the truncated gene fragments are listed in Table S1. All mutants produced by one-step homologous recombination were cultured in De Man–Rogosa–Sharpe (MRS) medium (Becton, Dickinson and company, New Jersey, USA) containing 10 μg/mL erythromycin (25) under static culture conditions. The mutant Δ*1932* (*rmlD2*) of YIT 9029 was constructed by two-step homologous recombination and described as Δ*cps1C* (20). The synthetic primer set for amplification of N- and C-terminal fragments are shown in Table 4. The plasmid pRD8 to isolate the mutant Δ*1932* (*rmlD2* deficient) of YIT 9029 is described in Figure S2.

### Reagents and chemicals for recombinant DNA technology

The reagents and DNA technology have been described in a previous report (20). Briefly, DNA was amplified by polymerase chain reaction (PCR) using KOD PLUS DNA polymerase (TOYOBO Co., Ltd., Osaka, Japan) or TaKaRa Ex Taq (Takara Bio Inc., Otsu, Japan). Restriction endonucleases, calf intestinal alkaline phosphatase, and a DNA Ligation Kit were purchased from Takara Bio Inc. or TOYOBO Co., Ltd. Plasmid purification was performed using the Wizard Plus SV Minipreps DNA Purification System (Promega K. K., Tokyo, Japan), and DNA fragments amplified by PCR were purified using the Qiaquick Gel Extraction Kit (QIAGEN K.K., Tokyo, Japan). Custom-made synthetic DNAs were purchased from Sigma-Aldrich Japan K. K. (Tokyo, Japan).

### Recombinant plasmid construction and insertion and deletion mutagenesis

The plasmids used in this study are listed in Table 2. The basic procedures for constructing recombinant plasmids for insertional mutagenesis, deletion mutagenesis, and isolation of bacterial clones harboring these chromosomal mutations have been described previously (20). Plasmid pRD8 was constructed as follows: DNA fragments containing N- and C-terminal coding regions of *CDS1932 (rmlD2)* gene were amplified by PCR using the primers shown in Table S1 and the purified DNA fragments thus obtained were digested with the respective restriction enzymes. These two fragments and pBE31 (25) digested with *Kpn* I and *Xba* I were mixed and ligated to obtain an in-frame deletion fragment of *rmlD2* gene, which was cloned on pBE31 (25). The resulting plasmid was named pRD8 (Figure S2). Construction of deletion mutants of YIT 9029 at *cps1A* (Δ*cps1A*) and *cps1C* (Δ*cps1C*), and each revertant harboring respective wild type (WT) gene in trans on the chromosome (Δ*cps1A,* Δ*cps1A/cps1A*, and *Δcps1C/cps1C*) were described previously (20). All primers for deletion mutagenesis were designed to enable in-frame rejoining of the N- and C-terminal peptide fragments of the gene, thereby avoiding translational interruptions within an operon. YIT 9029 was transformed with these plasmids, and erythromycin-resistant clones (25) were selected. These clones contained recombinant plasmids integrated into either side of the respective gene fragments via homologous recombination. After several cycles of subculturing (one thousandth inoculation into fresh medium, followed by full growth), erythromycin-sensitive clones were screened and checked for reversion or deletion.

### Culture of bacterial strains and storage

Bacterial cells were cultured in 4 mL of MRS medium with or without erythromycin (10 µg/mL) for 22-24 h at 37 °C under static culture conditions. After culturing (total cells: 1 × 10^9^–2 × 10^10^), the turbidity of the cultures was measured using a Klett-Summerson spectrophotometer (Klett MFG, New York, USA). Because the Klett value increased linearly with culture time up to 23 h (Figure S3), the culture time for each mutant strain was fixed to be 22 h in this study. Then the cells were labeled with 10 μM SYTOX Orange Nucleic Acid Stain (26) (Molecular Probes Co., Ltd.) in phosphate-buffered saline (PBS) containing 1% bovine serum albumin (BSA) for the lectin microarray analysis. The fluorescence intensity of 2 × 10^8^ labeled cells was measured using an ARVO X3 apparatus (PerkinElmer) after 1 h of labeling. The florescence of cells labelled with SYTOX Orange (26) were stabilized by freezing at −20 °C for 2 weeks. Hence, we stored labeled cells at −20 °C, and used within 2 weeks. The fluorescence intensities for all the tested strains were adjusted to within two times (Figure S3). Before testing for microarray, the cells were suspended in 360 μL of PBS containing 1% BSA (PBS/BSA) (27).

### Determination of the reactivity of mutants to YIT 9029-specific monoclonal antibody

The reactivity of the mutants to YIT 9029-specific MAb (21) was determined using a sandwich enzyme-linked immunosorbent assay (ELISA), as described previously (20). The resultant fluorescence intensities of WT (YIT 9029) and mutants were classified into three types, “positive” with full to half of the color intensity that WT showed, “slightly positive” with weak or slight color intensity, and “negative” with very weak or no color. An absorbance of ≥ 1.0 and < 0.25 was considered positive and negative. The absorbance of ≥ 0.25 but < 1.0 was considered slightly positive.

### Lectin microarray hybridization

The lectin microarray was prepared as described previously (28). Briefly, 96 lectins were dissolved at a concentration of 0.5 mg/mL in a spotting solution (Matsunami Glass) and spotted onto epoxysilane-coated glass slides (Schott) in triplicate using a non-contact microarray-printing robot (MicroSys4000; Genomic Solutions, Ann Arbor, MI). The origins and binding specificities of the tested lectins are listed in Table S3. YIT 9029, mutants of YIT 9029, and other tested strains were labeled with SYTOX Orange (26) as described above, then were added to each well of a glass slide containing immobilized lectins (100 μL/1-2 × 10^9^ cells/well) followed by incubation at 4 °C for 1 h. In this study, washing buffer at room temperature (RT) was used because no difference was observed in the binding of lectin to bacteria for the temperature of washing buffer between RT and 4 °C. Unbound cells were mildly removed by immersing the inverted lectin microarray glass slides in more than 1 L of PBS at RT for 30 min. Cells bound with lectins immobilized on a glass slide were detected using an evanescent-field fluorescence scanner. Data are shown as the ratio of the fluorescence intensities of the 96 lectins (29) relative to the maximal fluorescence intensity on the lectin microarray. Levels of lectin-binding signals are indicated by a color change from blue (low binding levels) to black (high binding levels).

### Effects of simple saccharides addition on the lectin microarray assay

To determine the effects of mono- and di-saccharides addition on the binding affinity of YIT 9029 to four lectins, rDiscoidin II, rBanan, Orysata, and CSA the following saccharides were added to the assay system at a concentration of 1 or 50 mM: D-galactopyranose (Gal), D-glucopyranose (Glc), D-galactosylpyranosyl-(1→4)-D-Glc (Lac), D-mannopyranose (Man), L-rhamnopyranose (Rha), D-fructofuranosyl-(2→1)-D-glucopyranoside (Suc), or (3S,4R,5S,6S)-6-Methyltetrahydro-2H-pyran-2,3,4,5-tetraol (Fuc). The efficiency of the inhibition of YIT 9029 binding to these lectins is shown as the ratio of the fluorescence intensity with each saccharide to that of the control with no added saccharides (27, 30–33)

### Profiling 51 mutants of YIT 9029

We compared the lectin-binding profiles of 51 YIT 9029 mutants (Table 1), YIT 9029 (WT), YIT 9022, YIT 9036, YIT 9037, and *L. casei* ATCC 334 (YIT 0180) with 96 lectin probes (Figure S4, Table S3) using advanced lectin microarray technology (27). To evaluate the similarity of gene functions, cluster analysis was performed to relate the cell wall PS biosynthesis genes in YIT 9029 and the lectin binding profiles. Considering the error between the arrays, YIT 9029 used two arrays in comparison between arrays. Unsupervised clustering was performed by employing the average linkage method using open-source Cluster 3.0 software developed by Michael Eisen of the Berkeley Lab. A heat map with clustering was generated using Java Treeview (Figure 4).

## RESULTS

### Identification of a cluster of genes associated with the cell wall biosynthesis of YIT 9029

In a previous study, we identified the biosynthesis genes of LCPS-1; cps1 gene cluster consisted of 10 genes designated *cps1A* to *cps1J* (GenBank AB470649) (20). To further analyze the genes responsible for the glycosylation of cell surface molecules of *L. paracase*i, we attempted to identify the genes of YIT 9029 that may participate in the biosynthesis and modification of cell wall PS, TA, and PG, including enzymes for sugar conversion, glycosylation, capsular PS polymerization, and PS repeat unit transporters from the whole genome sequence of YIT 9029. Based on amino acid sequence similarity to known genes of other bacterial strains, we selected 51 candidate genes, including the 10 *cps1* cluster genes described above (Table 1). *RmlA, rmlC, rmlB,* and *rmlD* are the biosynthesis genes of the Rha substance to Glc for cell-wall PSs (34), and two sets of *rml* cluster genes were found on the chromosome of YIT 9029 (Table 1). Unidentified genes located within possible operons of the PS synthetase genes were also included. For glyco-phenotype analysis, gene-deficient mutants were constructed toward these 51 genes having either deletion (Δ*1932* and Δ*cps1C*) or insertion (marked “Ω”, described in material and methods) within the individual genes (Table 2). Target genes are shown as scattered images in the genome of YIT 9029 (Figure S1).

### Ability of the mutants to bind to YIT 9029-specific MAb

The YIT 9029-specific MAb recognizes LCPS-1 on the cell surface of YIT 9029. The abilities of the MAb to the mutants were determined by the ELISA method (21) Based on the color intensity expressed as actual intensities of absorbance (Table S2), the mutants were largely categorized into three groups: positive, slightly positive, and negative (Table 3). Only seven of the 51mutants were totally negative for binding to MAb (21), namely *cps1A, cps1B, cps1C, cps1D, cps1E, cps1G,* and *cps1J,* constituting the *cps1* cluster (20). Eight mutants, including *cps1F*-deficient mutant, were classified into the slightly positive group and showed decreased antibody reactivity. Thirty-six of 51 mutants were classified as positive, indicating that there was no apparent change in the cell surface PS structure due to these mutations. These mutants are thought to conserve the cell surface structure recognized by the YIT 9029-specific MAb (21). Moreover, the reactivity of YIT 9029-specific MAb (21) in our laboratory collection strains with different antibody reactivity was as follows: YIT 9021 (24), YIT 9022, YIT 9036, and YIT 9037 were slightly positive, positive, negative and negative, respectively (Table 3).

**Table 3.** Reactivity of *L. paracasei* Shirota (YIT 9029) and the mutants to *L. paracasei* YIT 9029-specific monoclonal antibody. (**21**).

|  |  |
| --- | --- |
| Positive strains | YIT 9029 (Wild Type), $\Omega 0209$, $\Omega 0211$, $\Omega 0214$, $\Omega 0215$, $\Omega 0216$, $\Omega 0228$, $\Omega 0229$, $\Omega 0230$, $\Omega 0231$, $\Omega 0661$, $\Omega 0704$, $\Omega 0705$, $\Omega 0822$, $\Omega 0884$, $\Omega 0885$, $\Omega 1062$, $\Omega 1063$, $\Omega 1064$, $\Omega 1065$, $\Omega 1111$, $\Omega 1128$, $\Omega 1889$, $\Omega 1892$, $\Omega 1893$ (*rmlD1*), $\Omega 1894$ (*rmlB1*), $\Omega 1895$ (*rmlC1*), $\Omega 1896$ (*rmlA1*), $\Omega 1898$, $\Omega 1899$, $\Omega 1926$, $\Delta 1932$ (*rmlD2*), $\Omega 1934$ (*rmlC2*), $\Omega 1935$ (*rmlA2*), $\Omega 1937$ (*cps1I*), $\Omega 1938$ (*cps1H*), $\Omega 2708$, $\Delta cps1C/cps1C$, YIT 9022 |
| Slightly positive strains | $\Omega 0212$, $\Omega 0213$, $\Omega 0823$, $\Omega 0824$, $\Omega 0838$, $\Omega 1927$, $\Omega 1933$ (*rmlB2*), $\Omega 1940$ (*cps1F*), YIT 9021 |
| Negative strains | $\Omega 1945$ (*cps1A*), $\Omega 1944$ (*cps1B*), $\Delta 1943$ (*cps1C*), $\Omega 1942$ (*cps1D*), $\Omega 1941$ (*cps1E*), $\Omega 1939$ (*cps1G*), $\Omega 1936$ (*cps1J*), YIT 9036, YIT 9037, YIT 0180 |

The reactivity of the mutants to YIT 9029-specific monoclonal antibody was determined using a sandwich enzyme-linked immunosorbent assay, as described previously (21). The resultant fluorescence intensities of wild type and mutants were classified into three types: “positive” with full to half of the color intensity as that of wild type LcS, “negative” with very weak or no color, and “slightly positive” with weak or slight color intensity. An absorbance of $\geq 1.0$ was considered positive and $< 0.25$ was considered negative. The absorbance of $\geq 0.25$ but $< 1.0$ was considered slightly positive.

Therefore, the seven genes, *cps1A, cps1B, cps1C, cps1D, cps1E, cps1G,* and *cps1J,* among the 51 genes of YIT 9029 are essential to biosynthesize LCPS-1.

### Analysis of lectin binding properties of YIT 9029 and influence of *cps1C* gene on lectin binding affinity

We used a newly improved lectin microarray comprising 96 lectins, including 51 additional lectins (29). The lectin microarray format is shown in Figure 1A and the glycan-binding specificities of the lectins used in this study are listed in Table S3. YIT 9029 (WT) showed the affinity to an O-glycan binder (rDiscoidin II) and two Man binders (rOrysata and rBanana besides a Rha-binder CSA (formerly CSL) (Figure 1B left) (31–33, 35, 36).

**Fig. 1.**
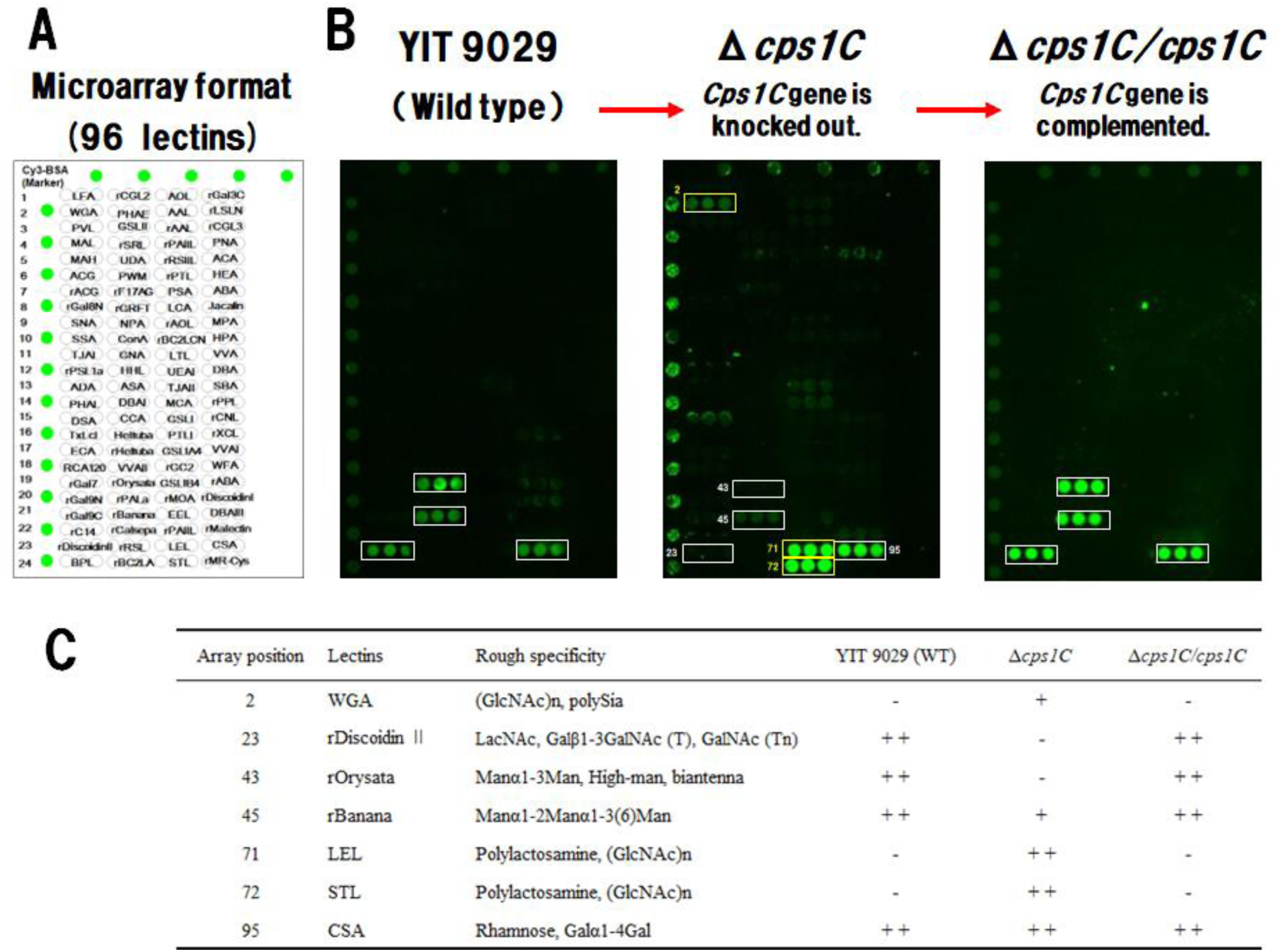
Result of lectin microarray analysis to evaluate the binding properties of YIT 9029 and influence of *cps1C* gene on lectin binding affinity. (A) Spot pattern of the lectin microarray with 96 lectins immobilized to a glass side in triplicate (29). (B) The lectin binding profiles of YIT 9029 (WT), Δ*cps1C* (*cps1C* gene is knocked out from YIT 9029), and Δ*cps1C/cps1C* (complemented by WT-*cps1C* to Δ*cps1C*) (21). YIT 9029 bound to an O-glycan binder (rDiscoidin II) and two Man binders (rOrysata, rBanana) besides a Rha-binder CSA (formerly CSL) (left). White frames indicate four types of lectins bound to WT-YIT 9029. Among the four lectins bound to YIT 9029, Δ*cps1C* did not bind to rDiscoidin II, Orysata, and rBanana, whereas it bound to CSA. On the other hand, Δ*cps1C* strongly bound to GlcNAc-binders WGA, LEL, and STL, but did not bind to WT-YIT 9029. Yellow frames indicate three lectins specifically bound by Δ*cps1C* (middle). The lectin binding profile of Δ*cps1C/cps1C* was completely recovered to that of YIT 9029 by complementing WT-*cps1C* gene into Δ*cps1C* (right). (C) Summary of the results. Each mark indicates binding affinity to lectins; ++ indicates strong, + indicates weak, and – indicates not binding. YIT 9029, *Lacticaseibacillus paracasei* strain Shirota; WT, wild type; Man, mannose; Rha, rhamnose.

Next, the influence of *cps1C* gene of YIT 9029 on the lectin-binding profile was investigated, which is essential for the biosynthesis of LCPS-1(20). The lectin binding profile of Δ*cps1C* was different from that of WT; the binding signal to rDiscoidin II and rOrysata disappeared, and that to rBanana reduced, whereas the binding to CSA was kept. Moreover, Δ*cps1C* strongly bound to WGA, LEL and STL but did not bind to YIT 9029 (Figure 1B middle) (31, 32, 35–38). These three lectins and LEL and STL, which are all GlcNAc-binders, often show specific preferences for polylactosamine and LacdiNAc (GalNAcb1-4GlcNAc) structures, respectively. Other lectins to which Δ*cps1C* weakly bound will be described later.

On the other hand, the lectin-binding profile of Δ*cps1C/cps1C* which is complemented by the WT-*cps1C* gene in Δ*cps1C* mutant, was the same as that of the WT (Figure 1B right). The results are summarized in Figure 1C. These data show that the lectin-binding profile of YIT 9029 is strictly controlled by *cps1C*, a key gene in the biosynthesis of LCPS-1 (20).

YIT 9029 binds to three additional lectins, rDiscoidin II, rOrysata, and rBanana, in addition to CSA (31–33, 35, 36). Moreover, these data indicate that the binding patterns of lectins to bacterial cells are closely associated with the gene functions involved and reflect the cell surface glycosylation profile.

### Carbohydrate inhibition assay

The binding of lectins to YIT 9029 was predicted to be mediated by the affinity of lectins for the carbohydrate moiety of cell surface structures. To speculate and clarify the affinity points for rDiscoidin II, rOrysata, rBanana, and CSA on YIT 9029 (Figure 1B), we attempted to detect possible interference of lectin binding by simple saccharides. We chose Gal, Glu, Lac, Man, Rha, Suc, and Fuc as inhibitors because each lectin bound to YIT 9029 is known to have an affinity for some glycoproteins containing some of these saccharides (27).

The binding affinity of YIT 9029 to rOrysata was inhibited by the addition of 1 mM Gal and Man, and 50 mM Glc, Man, and Suc (Figure 2 left). Similarly, rBanana was inhibited by the addition of 1 mM Man (Figure 2 middle). The binding affinity of YIT 9029 to CSA was inhibited by the addition of 50 mM Rha (Figure 2 right). The binding affinity of YIT 9029 to rDiscoidin II was not inhibited by any saccharides used in this study (data not shown). These data indicate that Man competitively inhibited the binding to rOrysata and rBanana, whereas Rha competitively inhibited the binding to CSA (Figure 2 right). Although there are some contradictions in the inhibitory effects of different saccharide concentrations, such as Suc for rOrysata and Rha for CSA, this may be due to the weak interactions between bacteria and lectins (31–33, 35, 36).

**Fig. 2.**
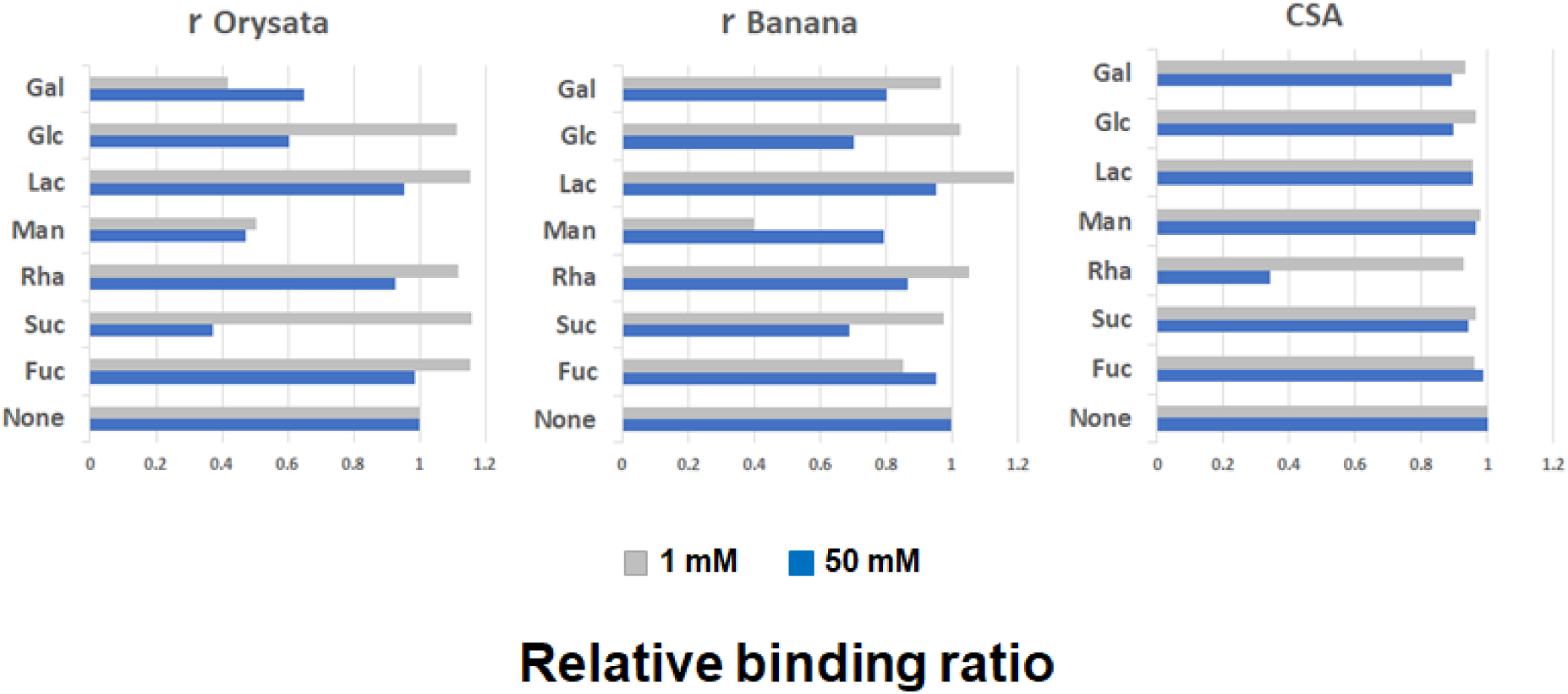
Carbohydrate inhibition assay. D-galactopyranose (Gal), D-glucopyranose (Glc), D-galactosylpyranosyl-(1→4)-D-Glc (Lac), D-mannopyranose (Man), L-rhamnopyranose (Rha), D-fructofuranosyl-(2→1)-D-glucopyranoside (Suc), or (3S,4R,5S,6S)-6-Methyltetrahydro-2H-pyran-2,3,4,5-tetraol (Fuc) were added in each well at a concentration of 1 mM (above) or 50 mM (below) with YIT 9029 cells (2 × 10^9^ cells/well) labeled with SYTOX orange. None means control assay that no carbohydrate is added to the reaction between lectins and YIT 9029. The efficiency of carbohydrate inhibition is shown as the ratio of the fluorescence intensity with carbohydrate against that of control with no added carbohydrate. The binding of YIT 9029 to rOrysata was inhibited by Gal, Glc, Man, and Suc (left), and that to rBanana and CSA was specifically inhibited by Man and Rha, respectively (middle and right).

These results confirm that the binding of YIT 9029 to rOrysata (31) is inhibited by a relatively large variety of sugars, such as Gal, Glc, Man, and Suc. Moreover, the binding affinity of YIT 9029 to rBanana and CSA was inhibited by Man and Rha, respectively (Figure 2) (5, 31, 35, 36).

### Effect of *cps1* cluster genes for LCPS-1 biosynthesis on lectin binding affinity

To analyze the effects of the genes responsible for biosynthesizing the cell wall of YIT 9029, we first determined the lectin-binding profiles of the gene knockout mutants within the *cps1* gene cluster, which are essential for LCPS-1 biosynthesis using microarray technology (Figure 3). The lectin-binding profiles of the 10 knockout mutants from *cps1A* (*CDS1945*) to *cps1J* (*CDS1936*) were unique to one another. When each binding profile were observed in more detail, similarities within groups were recognized. All these mutants except for Ω*1938* (*cps1H*) and Ω*1937* (*cps1I*) gained the abilities to bind to LEL and STL (39). The binding profiles of Ω*1938* (*cps1H*) and Ω*1937* (*cps1I*) were not different from that of the WT (YIT 9029) (Figure 1B left). Considering the fact that the YIT 9029-specific MAb (21) can bind to Ω*1938* (*cps1H*) and Ω*1937* (*cps1I*) (Table 3) and the previous report (20), it was concluded that the LCPS-1 structure (18) did not change in these mutants.

**Fig. 3.**
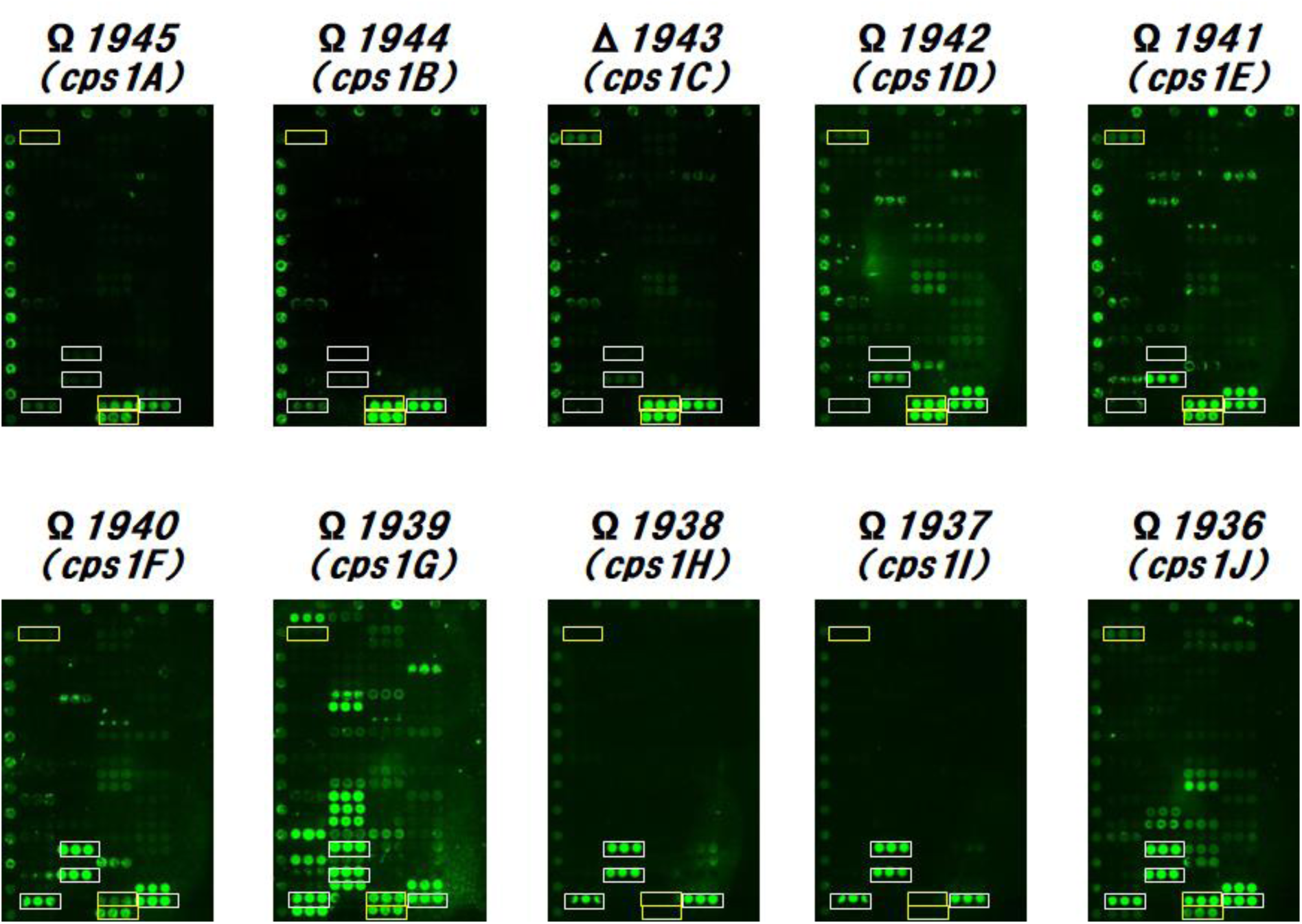
The effect of *cps1* cluster genes essential for LCPS-1 biosynthesis on lectin binding affinity. White frames indicate four lectins, rDiscoidin II, rOrysata, rBanana, and CSA, to which *L. paracasei* strain Sirota (YIT 9029) has specific affinities. Yellow frames indicate three lectins, WGA, LEL, and STL, to which Δ*cps1C* has specific affinities. The lectin binding profiles of Ω*cps1A*, Ω*cps1B*, and Δ*cps1C*; Ω*cps1D* and Ω*cps1E*; Ω*cps1F* and Ω*cps1J*; Ω*cps1H* and Ω*cps1I*, respectively, were well matched. The lectin binding profile of Ω*1938* (*cps1H*) and Ω*1937* (*cps1I*) were completely different from other eight mutants of disrupted *cps1* cluster genes, and similar to that of WT (YIT 9029) (Figure 1B left). LCPS-1, long-chain polysaccharide; WT, wild type.

The binding profiles of the mutants Ω*1945* (*cps1A)*, Ω*1944* (*cps1B*), and Δ*1943* (*cps1C*) were similar to one another because they lost the abilities to bind to rDiscoidin II and rOrysata and almost lost the ability to bind to rBanana (31, 32, 35, 36). Moreover, the binding profiles of the mutants Ω*1942 (cps1D*) and Ω*1941* (*cps1E*) were similar to each other due to loss of binding to LEL and rBanana and gain of binding to rMalectin (35, 36, 39, 40). In addition, these mutants showed affinity for some common additional lectins such as rF17AG, rAOL, and TJAII. The mutants Ω*1940* (*cps1F*) and Ω*1936* (*cps1J*) had similar binding profiles by gaining binding ability to LEL, STL and rMalectin, while maintaining the ability to bind to four lectins to which YIT 9029 binds (39, 40). However, it is obvious that the structural change in Ω*1940* (*cps1F*) is different from that of Ω*1936* because Ω*1940* partially bound to the YIT 9029-specific MAb, while Ω*1936* did not. The mutant Ω*1940* had a unique lectin binding profile among the *cps1* cluster genes knockout mutants (20), showing multiple lectin binding capacities.

In contrast, the lectin-binding profiles of Ω*cps1A*, Ω*cps1B*, Δ*cps1C*, Ω*cps1D*, Ω*cps1E*, Ω*cps1F*, and Ω*cps1J* were partially similar to one another. These eight mutants were negative or slightly positive for the YIT 9029-specific MAb (21) (Table 3). On the other hand, for three lectins, WGA, LEL, and STL to which Δ*cps1C* was bound (Figure 1B middle), these eight mutants strongly bound to both LEL and STL; however, the binding to WGA was not clear in Ω*cps1A*, Ω*cps1B*, and Ω*cps1G*. Other four lectins, rDiscoidin II, rOrysata, rBanana, and CSA to which YIT 9029 (WT) bound (Figure 1B left), had different lectin-binding properties. Although binding to CSA was observed in these eight mutants, binding to rDiscoidin II, rOrysata, and rBanana was different for each strain. Ω*cps1F*, Ω*cps1G*, and Ω*cps1J* strongly bound to these three lectins. Ω*cps1A*, Ω*cps1B*, and Δ*cps1C* weakly bound to those lectins. It was distinctive that Ω*cps1D* and Ω*cps1E* only bound to rBanana (31, 32, 35, 36, 41).

Moreover, the lectin binding profile of Ω*cps1G* was significantly different from other *cps1* gene-disrupted strains. Ω*cps1G* strongly bound to not only extensive lectins such as rDiscoidin II, rOrysata, rBanana, CSA, LEL and STL (5, 31, 32, 35, 36, 39), but also to other lectins such as LFA, RCA120, rGal9N, BPL, rF17AG, rGRFT, CCA, Heltuba, rHeltuba, rCalsepa, rAOL, rGC2, ACA, and rMalectin. The sugar-binding specificities of lectins are listed in Table S3.

Among the genes essential for LCPS-1 biosynthesis in YIT 9029 (20), lectin-binding properties differed greatly depending on the gene. These results strongly suggest that the effect on the cell surface structure changes specifically and drastically for each gene. Based on these results, it was possible to analyze the homology of gene functions by comparing the lectin-binding properties of genetically disrupted strains of YIT 9029.

### Profiling 51 mutants of YIT 9029

Based on the above results, a statistical analysis was performed to compare the lectin-binding profiles of 51 mutants of YIT 9029 (WT), two complementary strains, and the WT with 96 lectin probes (Figure S4, Table S3) using advanced lectin microarray technology (27). Our laboratory collection strains with different antibody reactivity of YIT 9021 (24), YIT 9022, YIT 9036, and YIT 9037 and *L. casei* ATCC 334 were also tested. The lectin binding properties of all tested strains were classified into three major clusters: from the top, the clusters were named 1, 2, and 3 (Figure 4). Clustering by lectin binding was consistent with the antibody reactivity (Table 3). Clusters 1 and 2 were composed of all positive strains and slightly positive strains to YIT 9029-specific Mab (21), except for Ω*cps1F*. Cluster 1 consisted of Ω*0213* alone, and it did not bind to the mannose-binder (rBanana) and showed weak binding to rOrysata (31, 35, 36). Cluster 2 included the largest number of members: 43 mutants and WT (YIT 9029), these mutant strains bound to an O-glycan binder (rDiscoidin II), two mannose binders (rOrysata and rBanana, and a Rha binder (CSA) (31, 32, 35, 36, 41). YIT 9021, YIT 9022, Δ*cps1A/cps1A*, and Δ*cps1C/cps1C* were also classified into cluster 2.

**Fig. 4.**
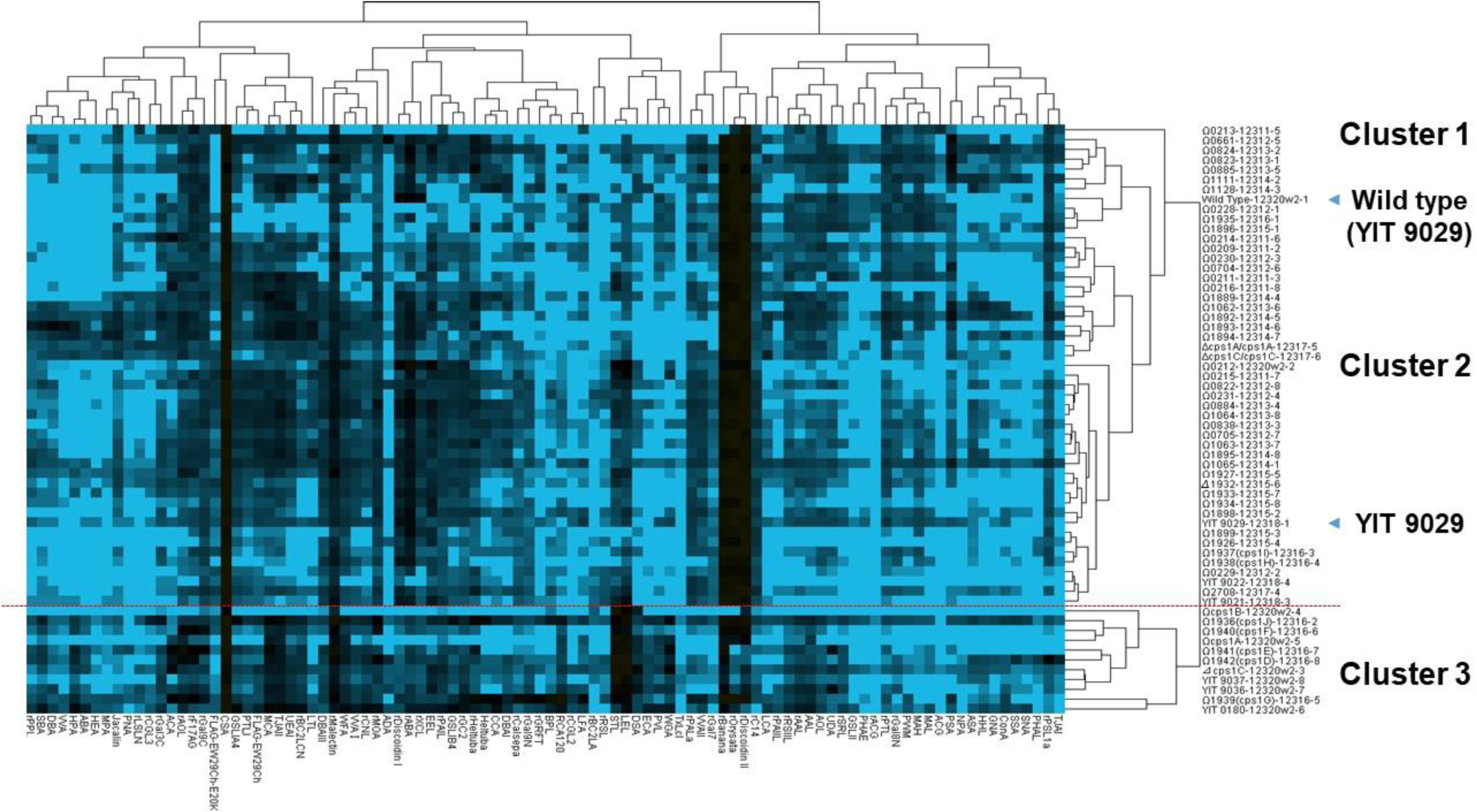
Relative binding of 51 mutants of *L. paracasei* strain Shirota (YIT 9029) with respect to lectin binding. Lectin-binding profiles of 51 mutants of YIT 9029 (WT) and WT with 96 lectin probes were compared using an advanced lectin microarray technology (27). YIT 9021(24), YIT 9022, YIT 9036, YIT 9037, and *L. casei* ATCC 334 were tested. The lectin-binding signals for each strain were normalized with the highest signal. The levels of lectin-binding signals are indicated by color change from blue (low binding levels) to black (high binding levels). The lectin binding properties of all tested strains were classified into three major clusters, from above, cluster one, two and three. Cluster one classified Ω*0213* alone, and that did not bind to the mannose-binder (rBanana) and weakly bound to rOrysata. Cluster two classified 43 mutants of YIT 9029 with YIT 9029 bound to an O-glycan binder (rDiscoidin II), two mannose-binders (rOrysata, rBanana), and a Rha binder (CSA). Cluster one and two had mutants of all positive strains and slightly positive strains except for Ω*cps1F* to YIT 9029-specific MAb reactivity. YIT 9021 (24), YIT 9022, Δ*cps1A/cps1A*, and Δ*cps1C/cps1C* were also classified in cluster two. Cluster three included Ω*cps1A*, Ω*cps1B*, Δ*cps1C*, Ω*cps1D*, Ω*cps1E*, Ω*cps1F*, Ω*cps1G*, Ω*cps1J*, YIT 9036, and YIT 9037. All these mutants had negative and only one had slightly positive strain (Ω*cps1F*) to YIT 9029-specific MAb reactivity. Moreover, Δ*cps1A/cps1A* was a positive strain to YIT 9029-specific MAb reactivity (20).

Cluster 3 included Ω*cps1A*, Ω*cps1B*, Δ*cps1C*, Ω*cps1D*, Ω*cps1E*, Ω*cps1F*, Ω*cps1G*, Ω*cps1J*, YIT 9036, and YIT 9037, all of that showed negative binding to YIT 9029-specific Mab (21), except for Ω*cps1F*, that was slightly positive to the antibody binding. Moreover, based on the lectin-binding affinities of our four laboratory collections with different antibodies, YIT 9021 and YIT 9022, and YIT 9036 and YIT 9037 were categorized in Cluster 2 and Cluster 3, respectively.

It was confirmed that the lectin-binding properties of the gene-disrupted strains closely matched the reactivity of these mutants with YIT 9029-specific Mab (21) with some exceptions. For the present analysis, we included some gene knockouts within the same predicted operon because genes in the same operon (transcriptional unit) often produce one final product. For this, the following four possible operon genes were chosen: Ω*0209* to Ω*0216*, Ω*0228* to Ω*0231*, Ω*0822* to Ω*0824*, and Ω*1062* to Ω*1065*. The lectin-binding profiles of the mutants within the same operon showed certain similarities. Cluster 2, composed of 43 mutants, showed profiles similar to that of YIT 9029. Though all mutants of this cluster bound to mannose-binders (rOrysata and rBanana), only Ω*0213* cells did not bind to rBanana (18). For this reason, only Ω*0213* cells were classified as a different branch from other strains which were classified under Cluster 1.

In contrast, all mutants of YIT 9029 bound to the Rha binder CSA (41). The reason for this is discussed later.

## DISCUSSION

It is well documented that LAB used in food are beneficial for human health; however, the interaction between bacterial cell surface components and host immune cells is limited. We investigated *Lacticaseibacillus paracasei* strain Shirota (YIT 9029) for its immunological action against host cells. YIT 9029 plays a critical role in the modulation of immune cells (20). YIT 9029 is a cured strain of the bacteriophage FSW of *Lactobacillus casei* YIT 9018 (42), and both strains react with YIT 9029-specific Mab (21). It has unique PSs called LCPS-1 (18) and LCPS-2 (19) on its cell surface, and LCPS-1 greatly affects the physiological effects of this bacterium.

Based on the previous reports (20, 27), we investigated how each of the genes involved in cell surface PS synthesis of YIT 9029 affects the actual structure of this bacterium by analyzing which genes are linked to which lectin-binding properties. To answer these questions, we isolated gene disruption mutants of YIT 9029, and 51 genes including the *cps1* cluster genes (20) were knocked out (Table 1). As a technical challenge, lectin microarray analysis was initially developed to analyze mammalian cells. However, it was successfully applied to analyze YIT 9029 mutants using an advanced version of 96 lectin microarray (29) (Figure 1A, Table S3). Hence, a series of gene disruption mutants of glycosylation genes and modification enzymes of YIT 9029 were constructed, and the structure and functional relationships of the cell surface molecules of YIT 9029 were elucidated in more detail. Furthermore, we compared their binding profiles to lectins in parallel with their binding abilities to YIT 9029-specific Mab that is our original tool (21). As shown in Figure 1B, YIT 9029 has an affinity for an O-glycan binder (rDiscoidin II), two Man-binders (rOrysata and rBanana), and Rha binder CSA (31, 32, 35, 36, 41). The binding affinities between YIT 9029 and Man-binding lectins, rOrysata/rBanana, were inhibited by 1 mM Man (Figure 2). Interestingly, YIT 9029 is not found in the cell wall PS of Man itself (18, 19). Although it is unclear what these lectins recognize in the cell wall PS structure of YIT 9029, the binding of various lectins to YIT 9029 is of interest.

Furthermore, it can be considered that the lectin binding pattern of each bacterial strain reflects the cell surface structural characteristics, especially glycosylated molecules. When the lectin-binding patterns of mutants with cps1 cluster genes were determined, the alterations in the binding patterns could be grouped as described in the RESULTS section.

In this study, several remarkable points were observed through the lectin microarray analyses. a) The loss of LCPS-1 did not influence the number of lectins capable of binding to mutant cells. We considered LCPS-1 as a suppressive molecule that induces cytokine production by macrophages owing to the physical masking effect of LCPS-1 on the cell surface (20). b) The Ω*1939* (*cps1G*) and Ω*1936* (*cps1J*) mutants still has the affinity to rDiscoidin II, rOrysata and rBanana, that are possible markers for LCPS-1 biosynthesis, although these mutants do not react with YIT 9029-spcific MAb reactivity, nor produce LCPS-1 (20). c) The similarity of the lectin-binding profiles of Ω*1942* (*cps1D*) and Ω*1941* (*cps1E*) mutants probably indicate that these gene products have a similar role in the synthesis and maturation of LCPS-1. d) The Ω*1939* (*cps1G*) mutant dramatically changes the binding pattern of lectins, indicating big changes in the surface structure of the mutant cells. All these indications and assumptions will be addressed in future research.

The mutation that reduced antibody reactivity in YIT 9029 was confirmed to be a deletion of LCPS-1 (20). However, it is unclear whether the effects of LCPS-1 deletion on the cell surface PS structures of YIT 9029 cells are uniform or diverse. Here, we demonstrated that the mutations in cell surface PS that occur after the deletion of LCPS-1 vary greatly depending on the gene.

Another interesting result is that while Ω*0213* develops LCPS-1 on the cell wall, the lectin binding profile is different from that of YIT 9029 (Figure 4). In this mutant, the *CDS0213* gene encoding a putative PS biosynthesis protein (CAQ65366.1) (22) was knocked out, and the sequence of *CDS0213* was similar to eps7I of *L. casei* strain BL23 (22) (Table 2). This gene is widely conserved in bacterial genome, and that is classified as the Glycosyltransferase Family 32 (http://www.cazy.org/GT32.html) with known activity as α-1,6-mannosyltransferase (EC 2.4.1.232). This result was a good match between the predicted gene function and lectin-binding in the bacteria.

Recently, the structure of LCPS-1 was clarified that was found to be more complex than that of LCPS-2. Since Rha is included in both LCPS-1 (18) and LCPS-2 (19) moieties, we expected that CSA would bind to both YIT 9029 and LCPS-1-negative mutants through the Rha residues of both LCPS-1 and LCPS-2 (Figures 3 and 4). The primary structure of LCPS-1 (18) contains Rha on the linear supralateral chain, and Rha on the linear chain is acetylated (32). Although the ligand structure recognized by the YIT 9029-specific MAb (21) reactivity has not yet been identified, it may recognize the acetyl group of LCPS-1.

On the other hand, among the 51 genes chosen for disruption, several groups of genes may form operons, including Ω*0209* to Ω*0216*, Ω*0228* to Ω*0231*, Ω*0822* to Ω*0824*, and Ω*1062* to Ω*1065*. As shown in Figure 4, we confirmed that the lectin-binding profiles of the mutants within the same operon have certain similarities.

These observations suggest that the combined use of lectin microarray technology and a mutant strain library of YIT 9029 is a powerful tool for identifying unknown bacterial gene functions in terms of cell surface glycome, that is a novel approach to glycophenotyping. It is important to clarify the relationship between the phenotype and genotype of bacterial strains. The use of lectins in combination is an effective solution when antibodies cannot be produced by the bacterial strain.

In conclusion, the combined use of lectin microarray technology and a mutant strain library of YIT 9029 makes a powerful tool for identifying unknown bacterial gene functions in terms of cell surface glycome. This is a novel approach to glycophenotyping, and lectin microarrays enable the identification of the cell wall glycome in relation to bacterial gene function.

## SUPPLEMENTARY DATA

Supplementary data for this article is available at www.////.org.

## ACKNOWLEDGMENT

We are grateful to Ms. Jinko Murakami at the National Institute of Advanced Industrial Science and Technology for her help with preparation of the lectin microarray. We thank Haruji Sawada, who carried out this project, and Hoshitaka Matsumoto, who reanalyzed the homology of YIT 9029 with *L. casei* BL23. We are deeply indebted to Ritsuo Aiyama and Teruo Yokokura, who always encouraged us, and to the late Toshiaki Osawa for pioneering research in lectins and constructive discussions.

---

**Fig. S1.**
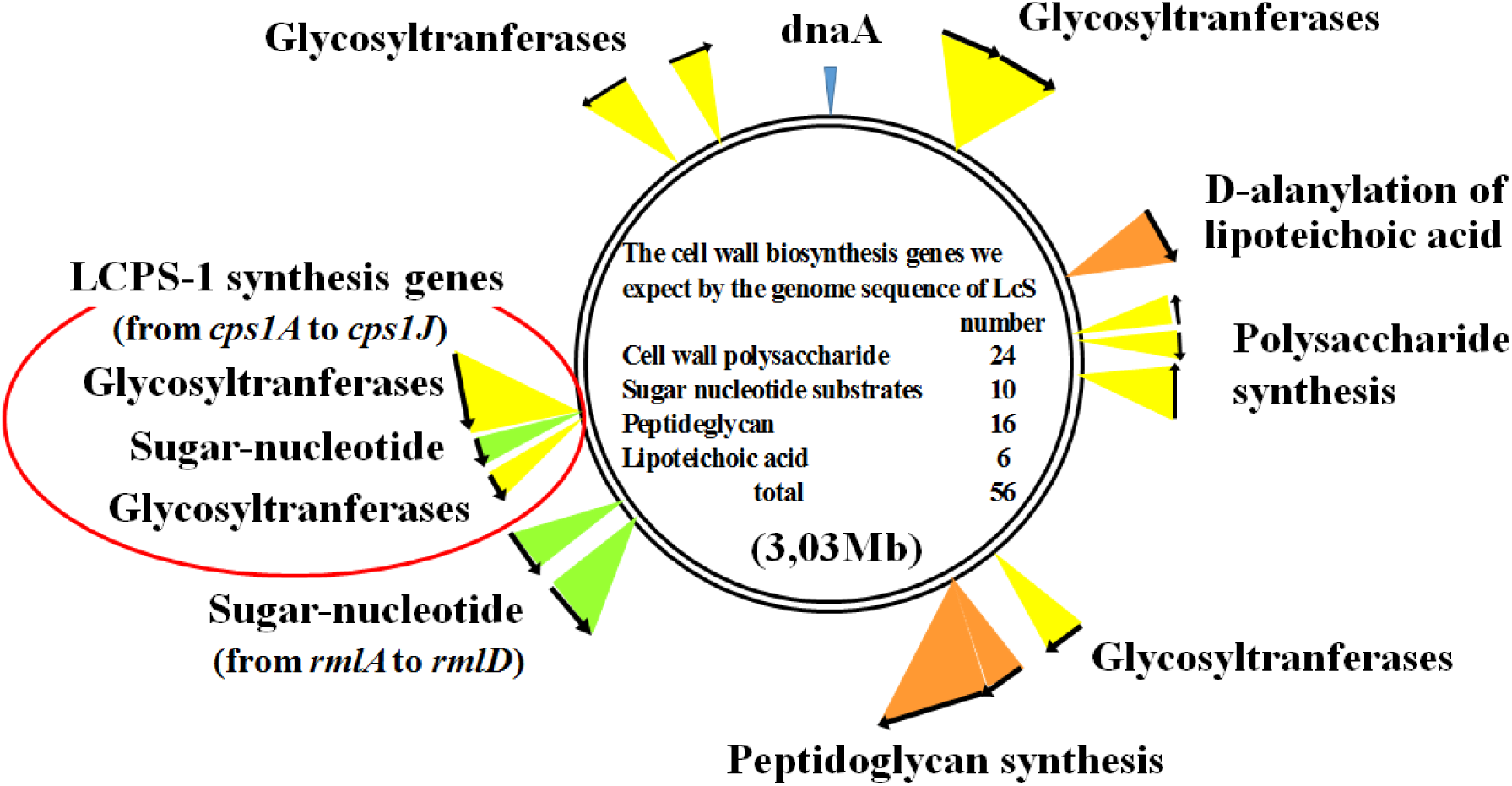
Distribution map of cell wall biosynthesis-related genes in the chromosome of YIT 9029. Fifty-six genes are required for the cell wall biosynthesis. Twenty-four, 10, 16, and six genes are required for the biosynthesis of cell wall polysaccharide, sugar nucleotide substrates, peptidoglycan, and lipoteichoic acid, respectively. We succeed in making 51 gene knockout mutants of YIT 9029. The remaining five were very difficult to isolate.

**Fig. S2.**
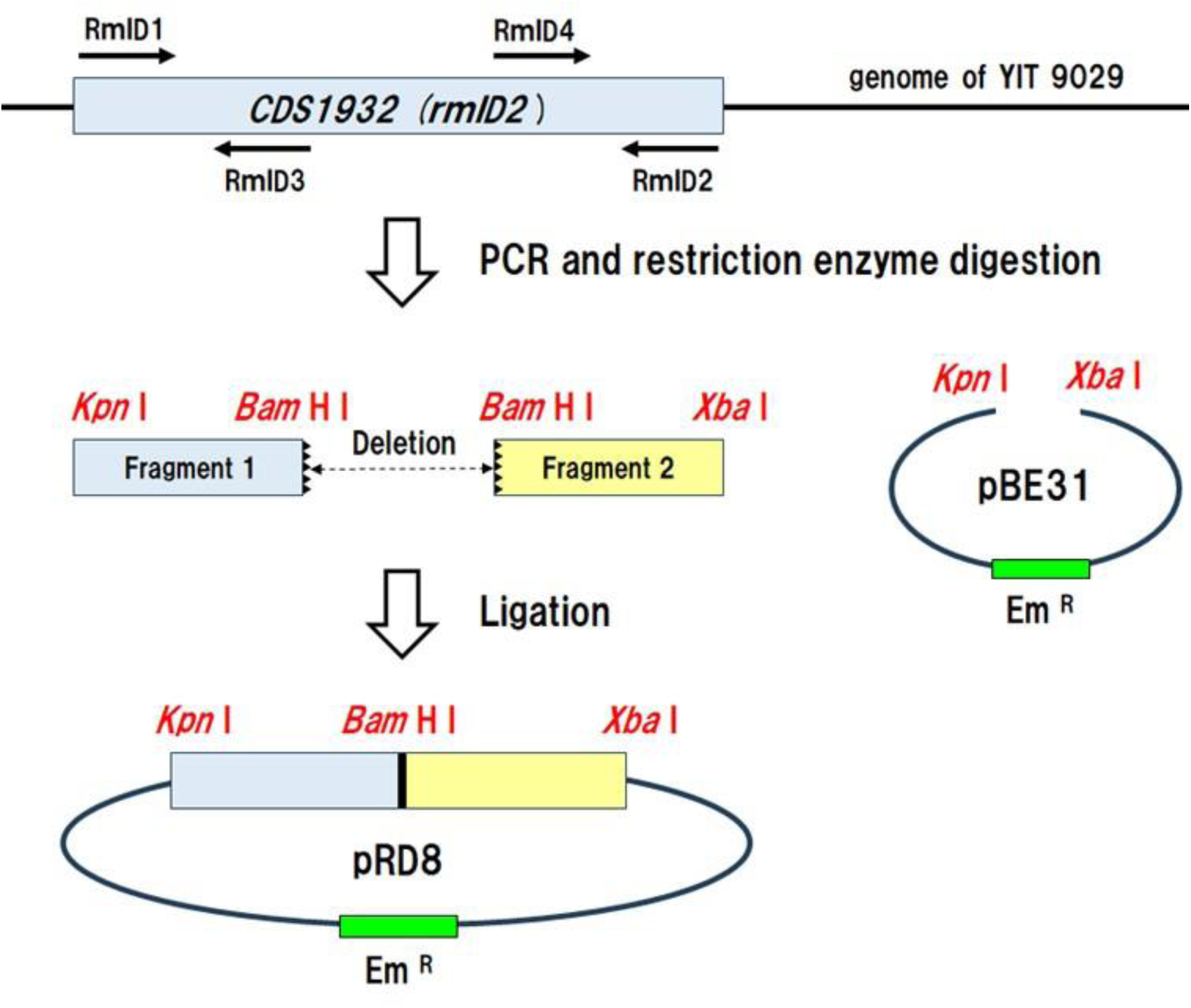
Preparation of plasmid pRD8. Plasmid pRD8 was constructed as follows; DNA fragment-1 containing the 5’-terminal of the *CDS1932 (rmlD2)* gene were amplified with primers of RmlD1 and RmlD3 (5’-GGGGTACCTA GAGGGGAAGA GAAATGAA-3’ and 5’- CGGGATCCTA AAGTGGCGTC AATGGCC-3’). DNA fragment-1 was digested with the restriction enzymes *Kpn* I and *Bam*H I. DNA fragment-2 of the *CDS1932* (*rmlD2*) gene were amplified with primers of RmlD4 and RmlD2 (5’-CGGGATCCGA ATATGGTCAC AACTTTGTT-3’ and 5’-GCTCTAGATT ATTGCTTTTT AATCACCTGC-3’). DNA fragment-2 was digested with the restriction enzymes *Bam*H I and *Xba* I. These fragments were cloned into pBE31 (25) digested with the restriction enymes *Kpn* I and *Xba* I in the same order as on the chromosome to obtain in-frame deletions within the genes. YIT 9029, *Lacticaseibacillus paracasei* strain Shirota; PCR, Polymerase Chain Reaction

**Fig. S3.**
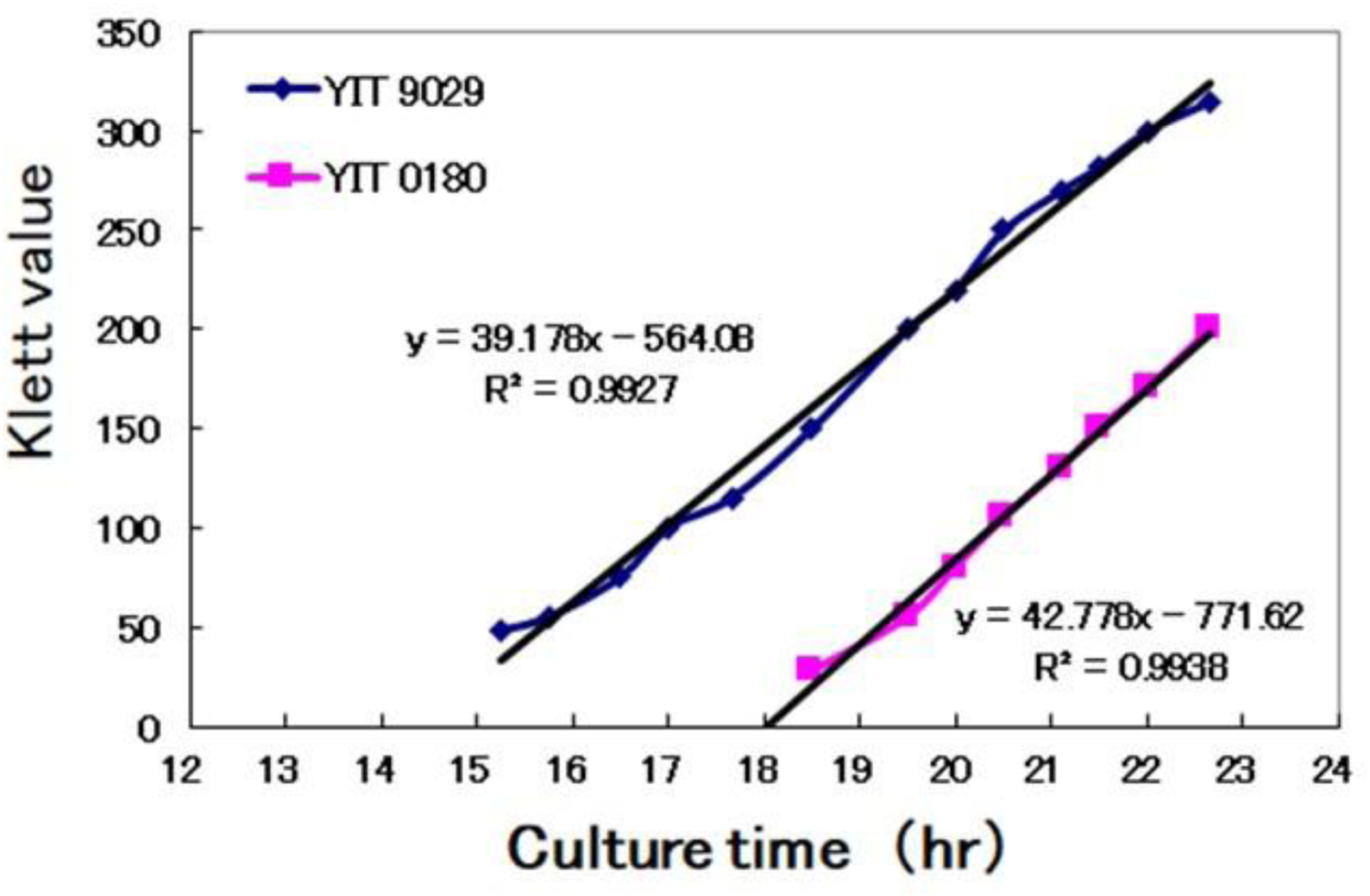
Klett value of *L. paracasei* strain Shirota YIT 9029 and *L. casei* ATCC 334 (YIT 0180) linearly increased with increase in culture time till 23 h at 37 °C under aerobic conditions. Klett value of 100 and 200 was over 1 × 10^8^ and 1 × 10^9^ (cells/mL), respectively, for these cells. Culture time of bacterial strains was 22-24 h.

**Fig. S4.**
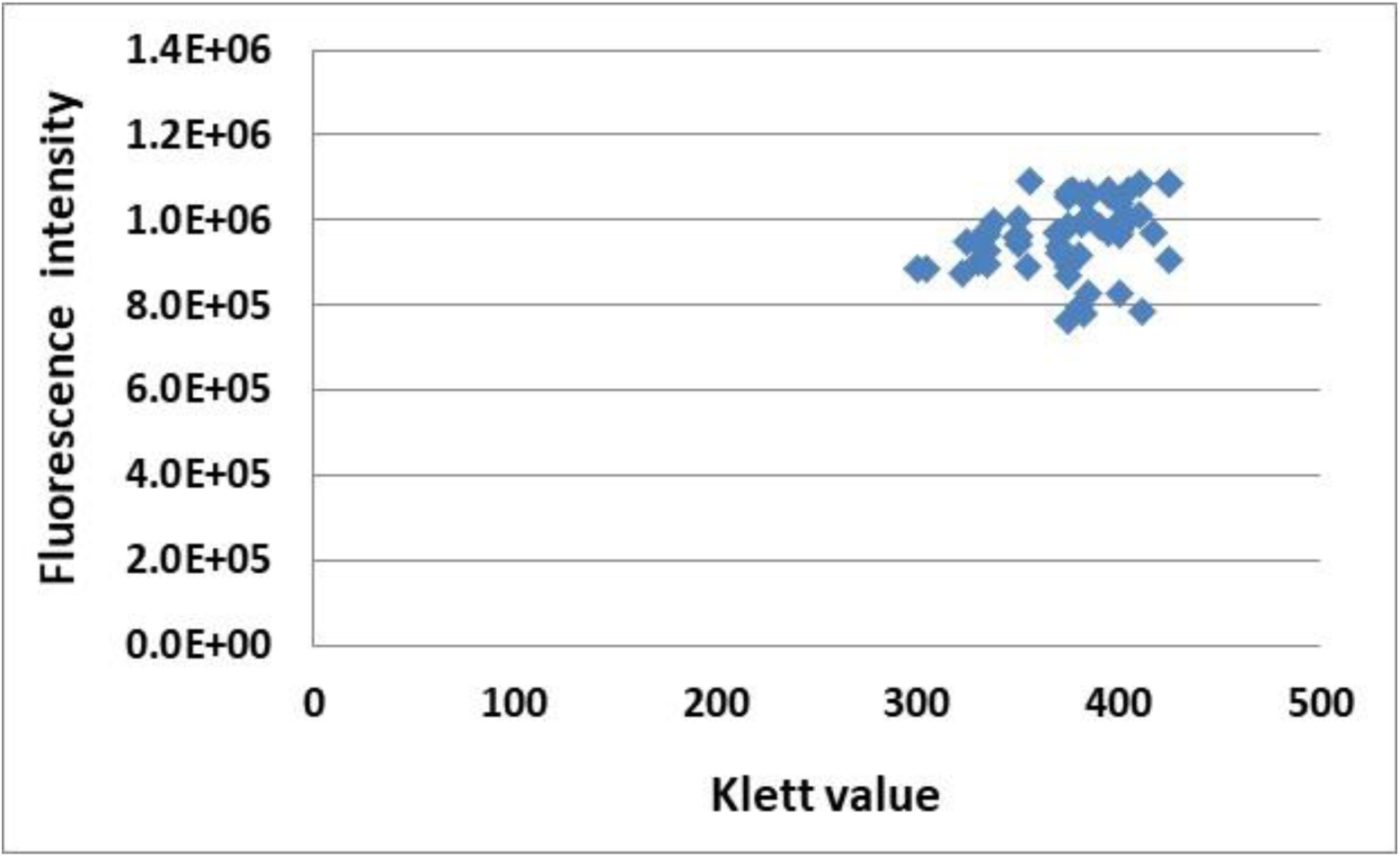
The fluorescent intensity of all tested strains. The fluorescent intensity of *L. paracasei* YIT 9029, all mutants, and our other collections were labeled by SYTOX Orange, which was adjusted ≤ 2 times. Klett value was over 300.

**Table S1.**
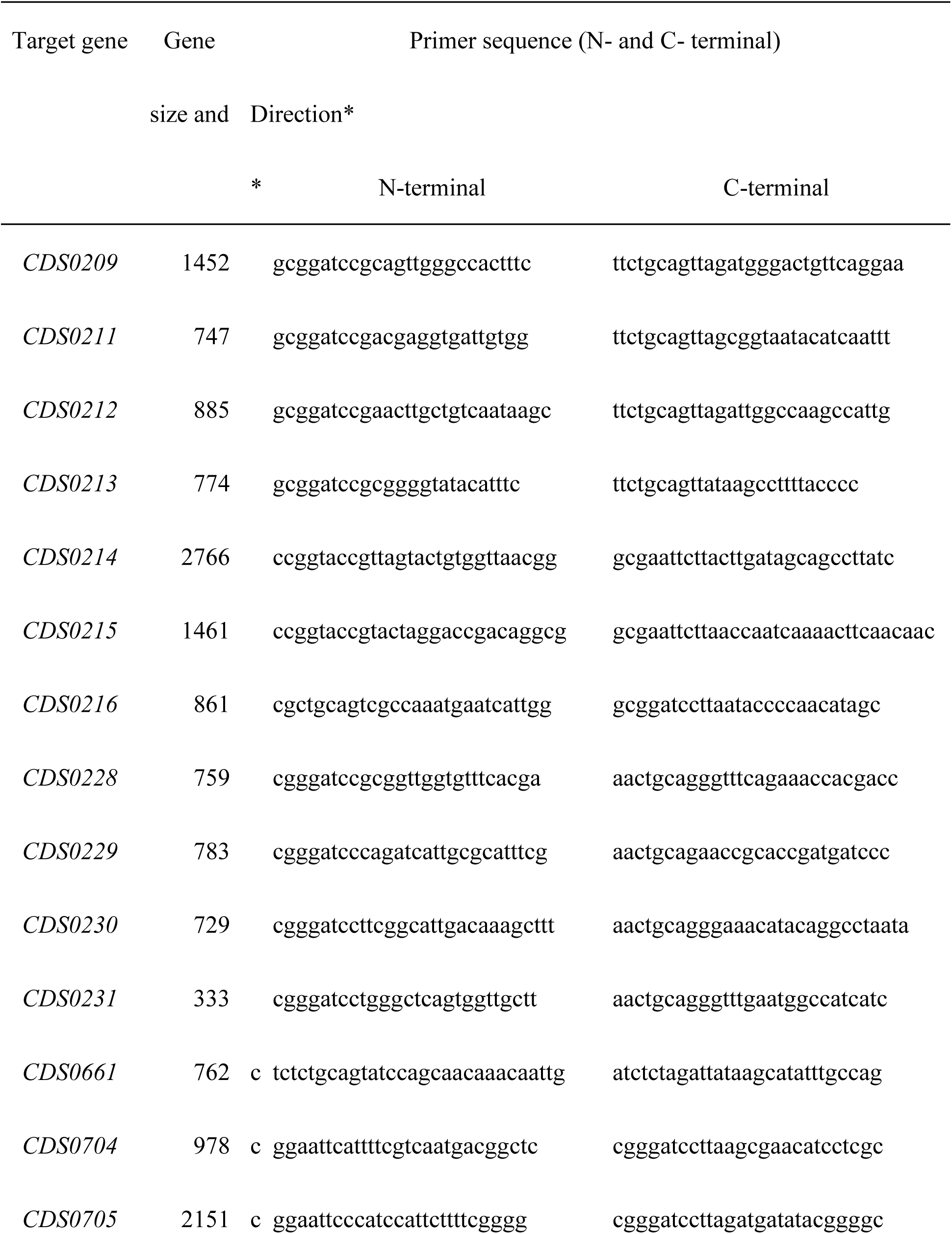
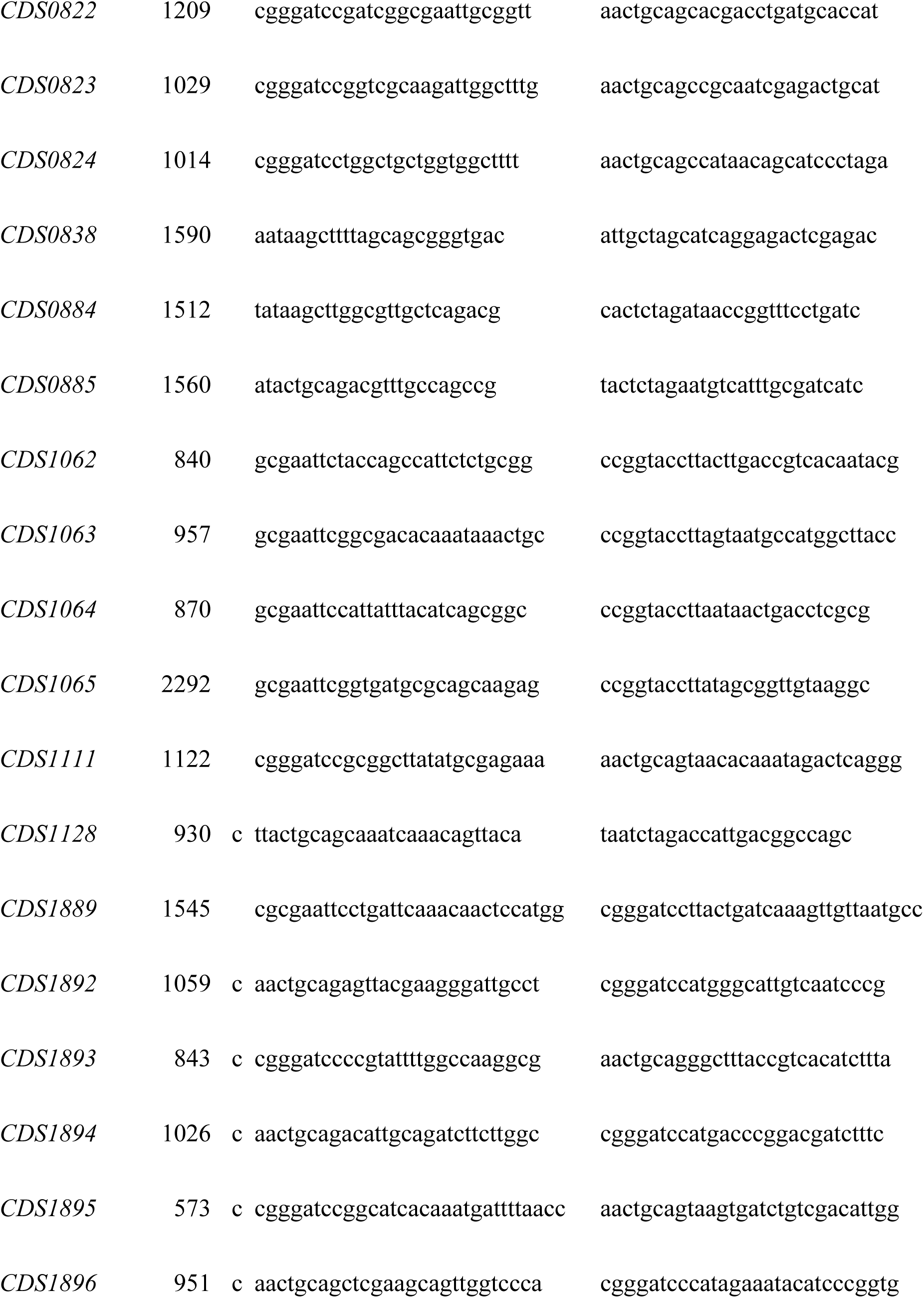
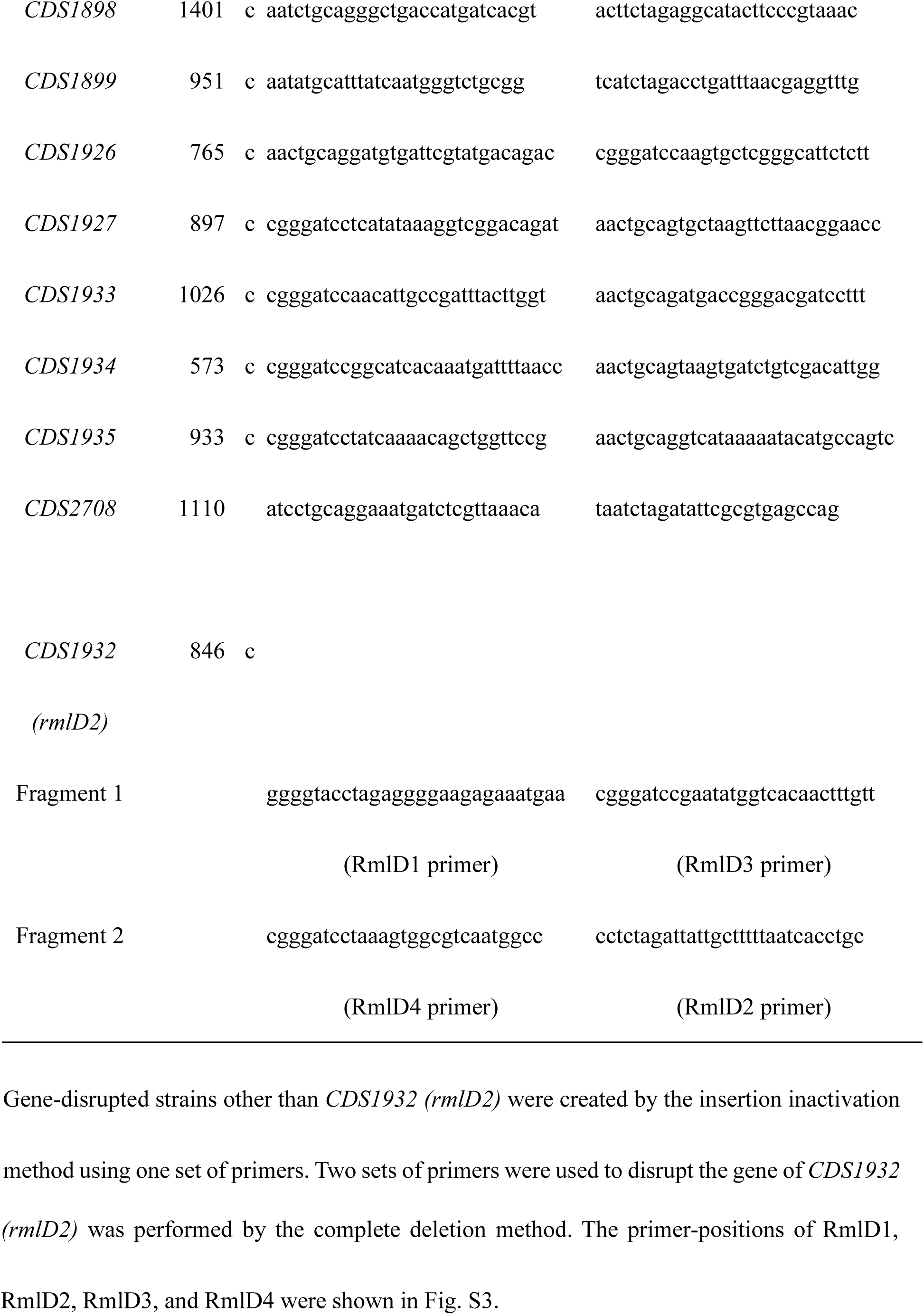
Synthetic primers for truncated amplification.

**Table S2.**
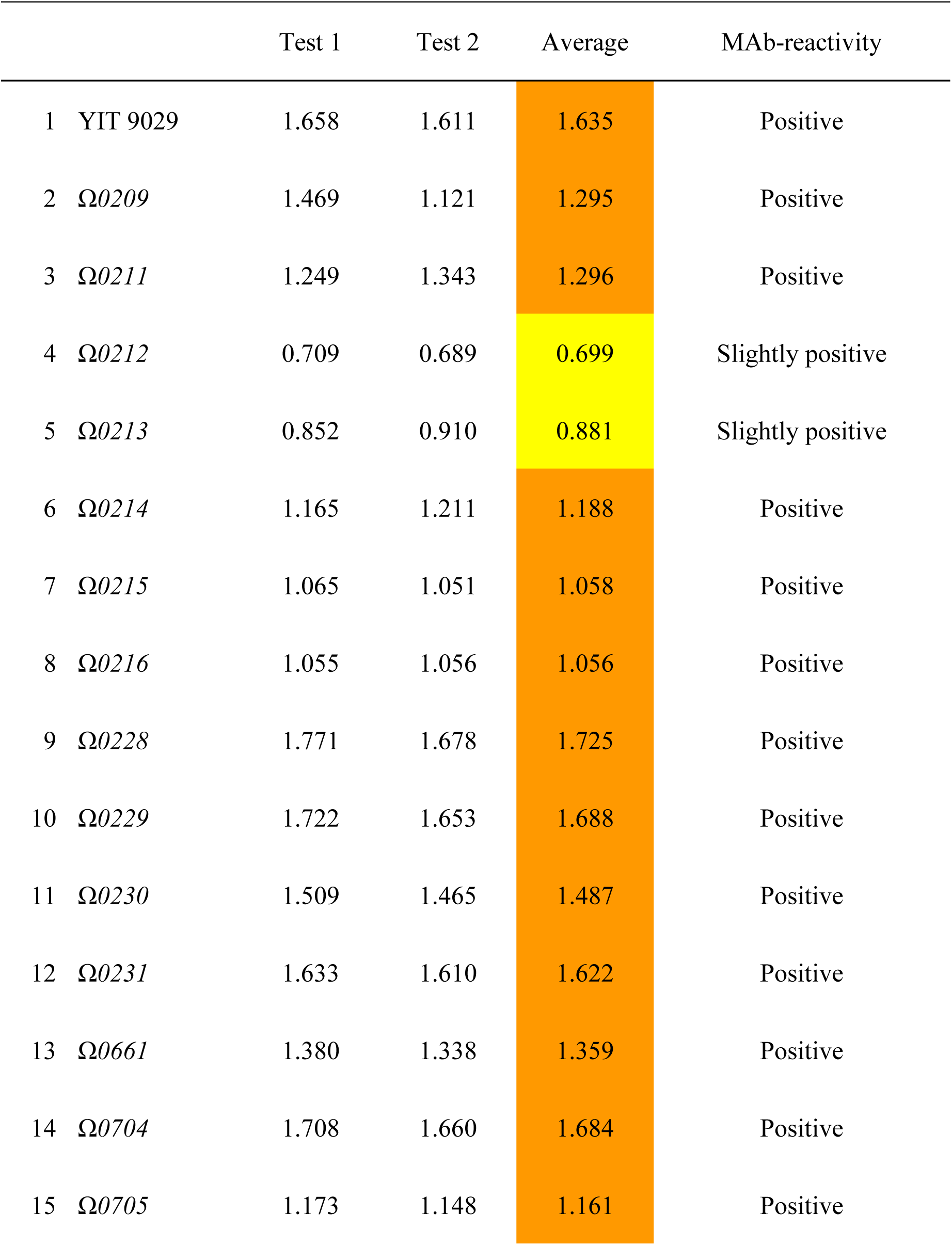
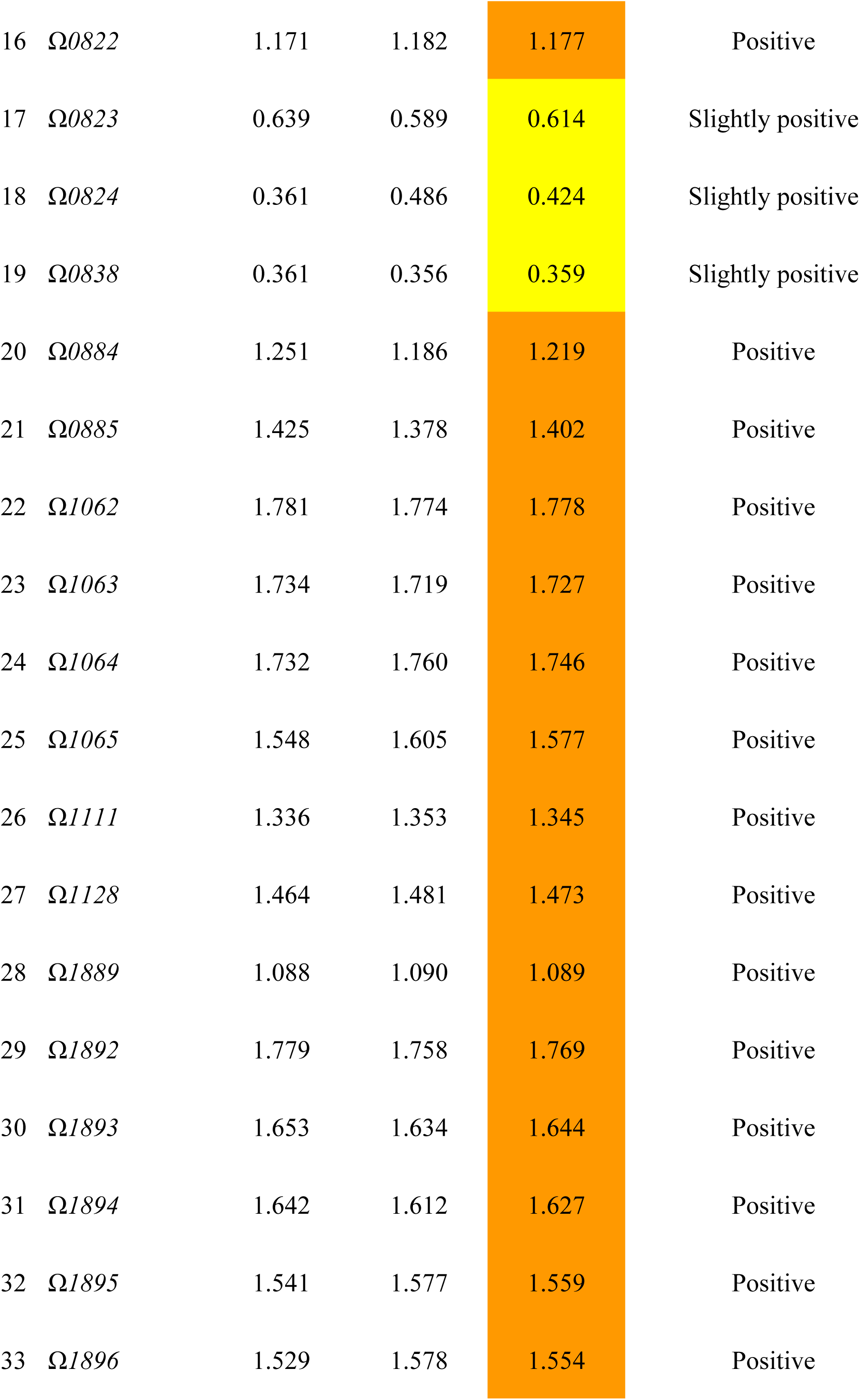
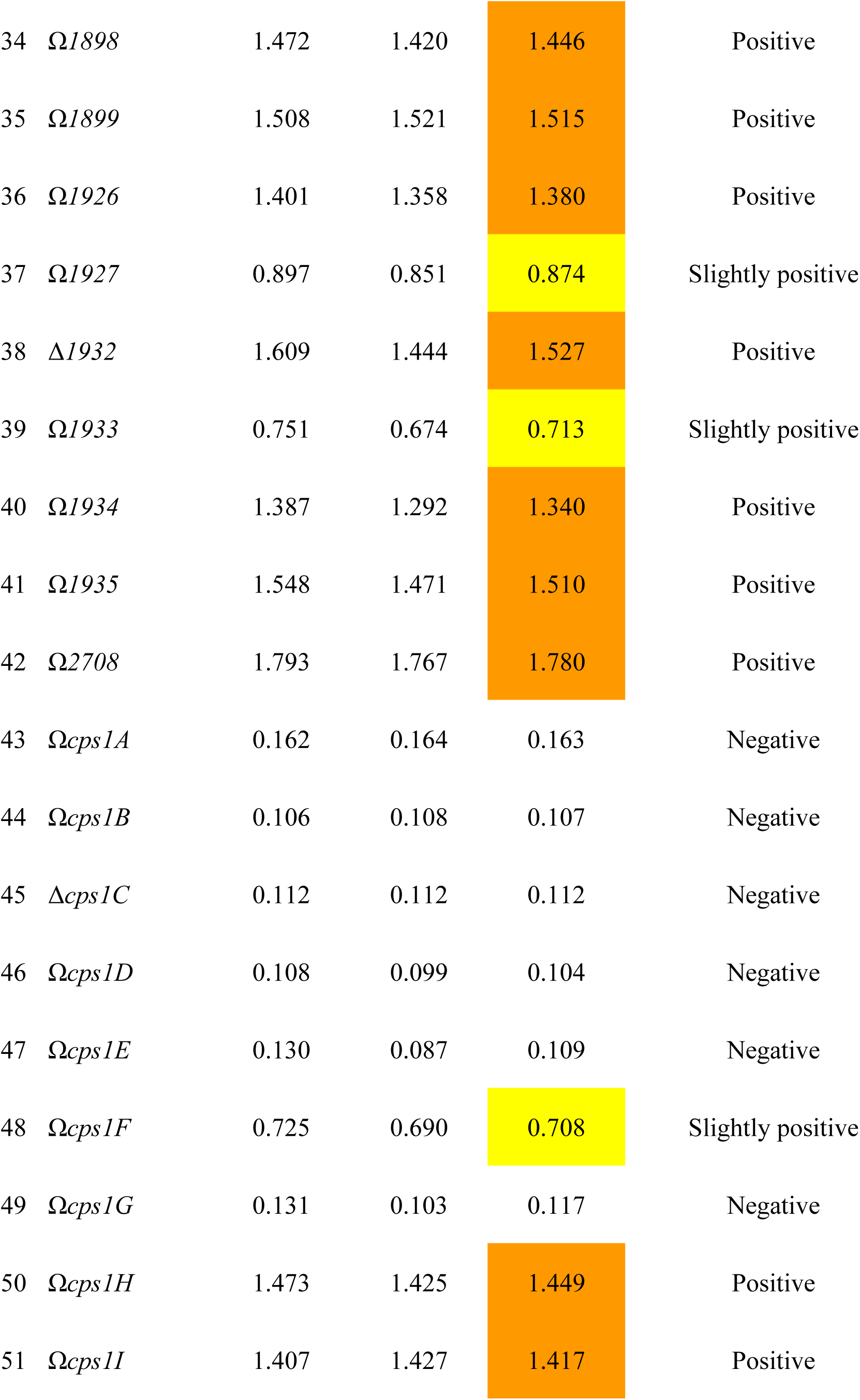
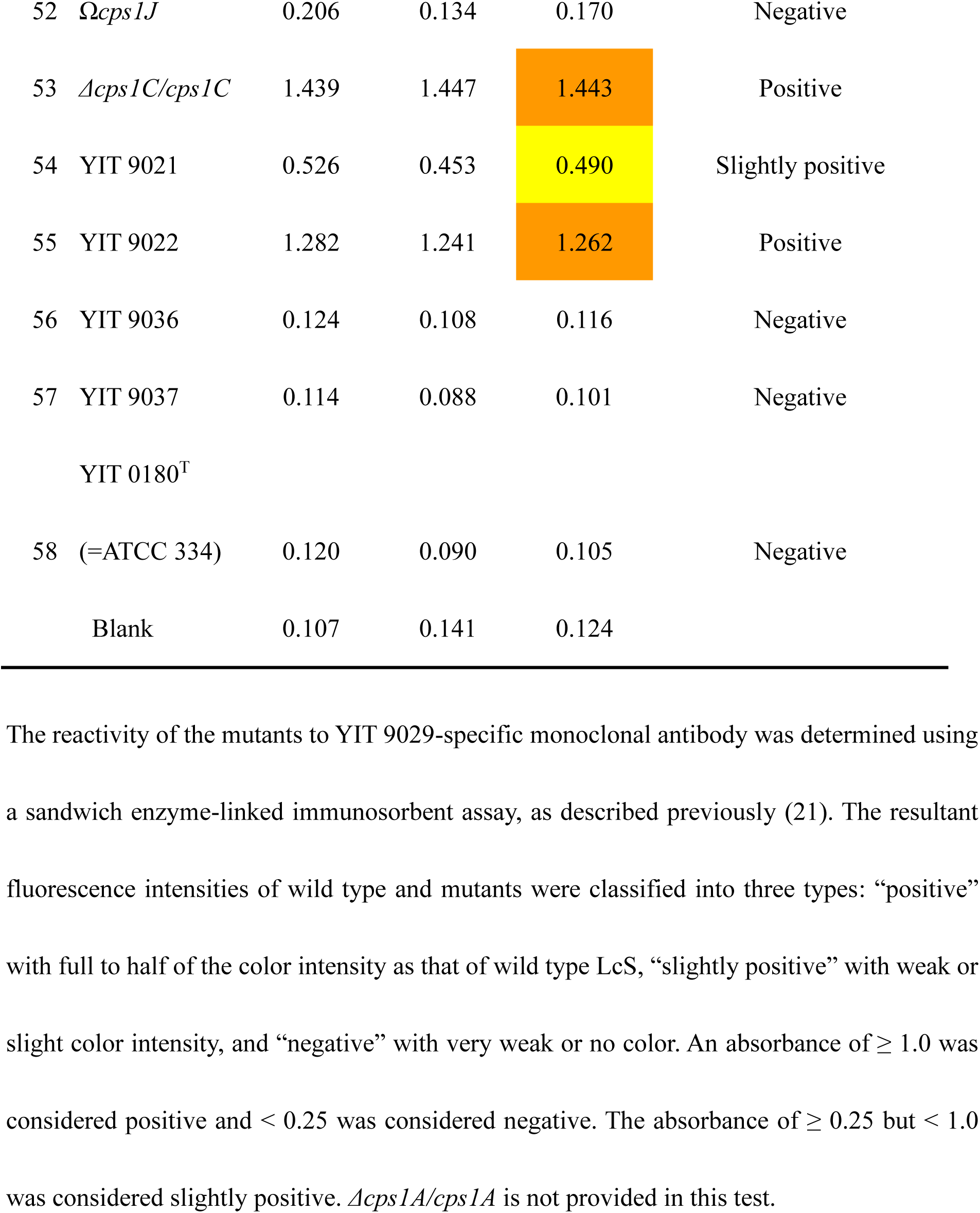
Reactivity of *L. paracasei* Shirota (YIT 9029) and mutants to *L. paracasei* Shirota-specific monoclonal antibody (MAb)

**Table S3.**
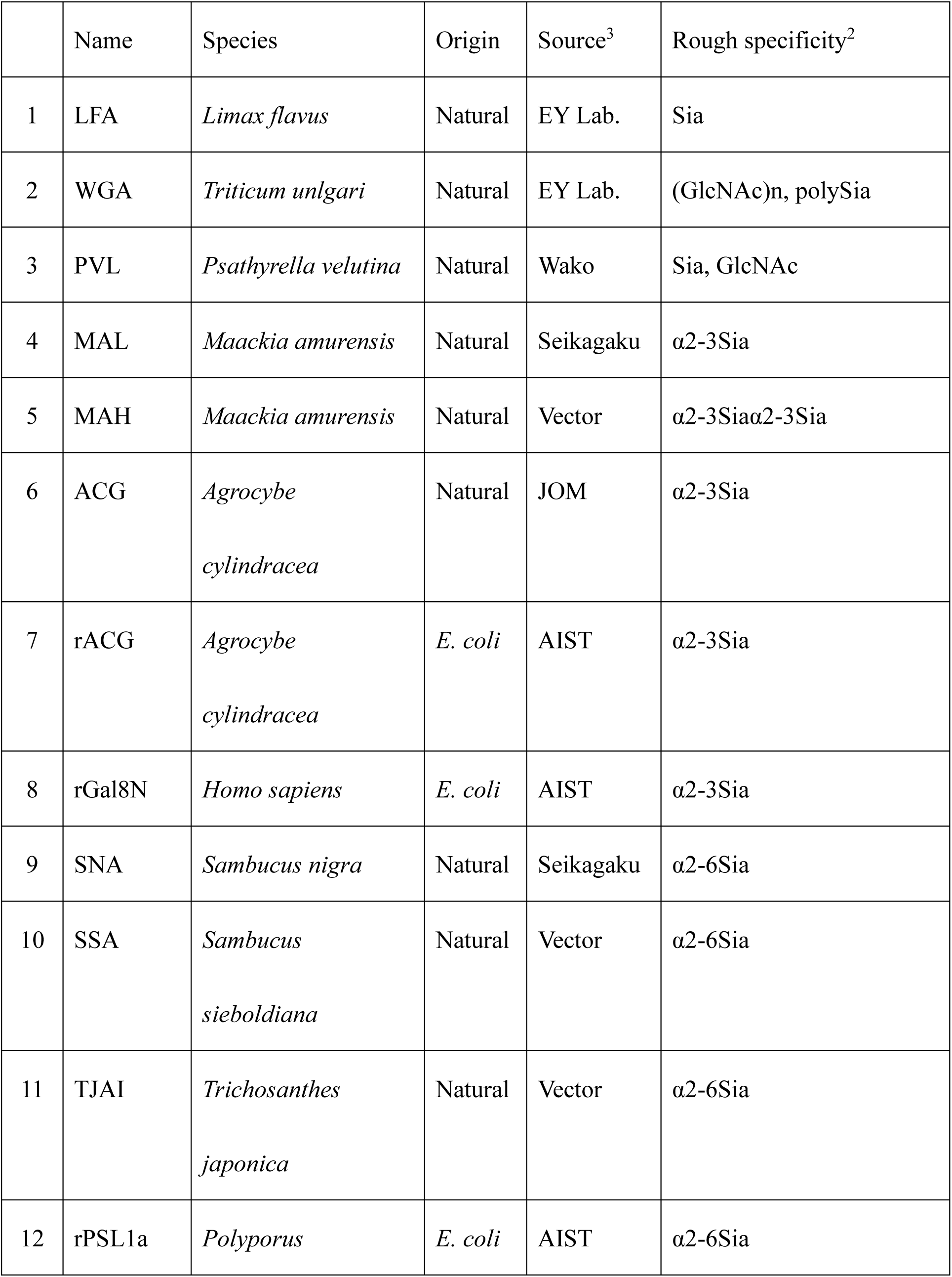
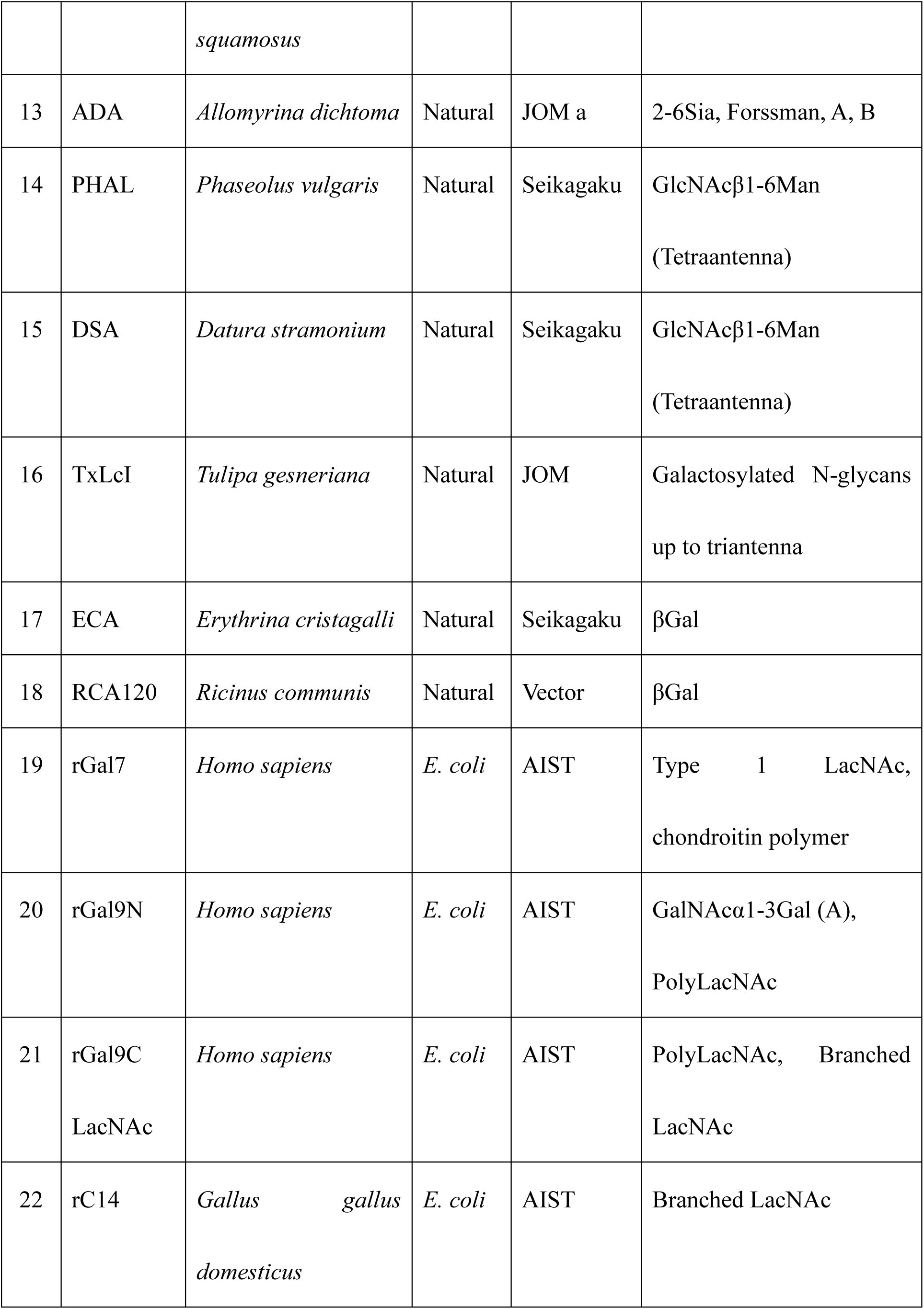
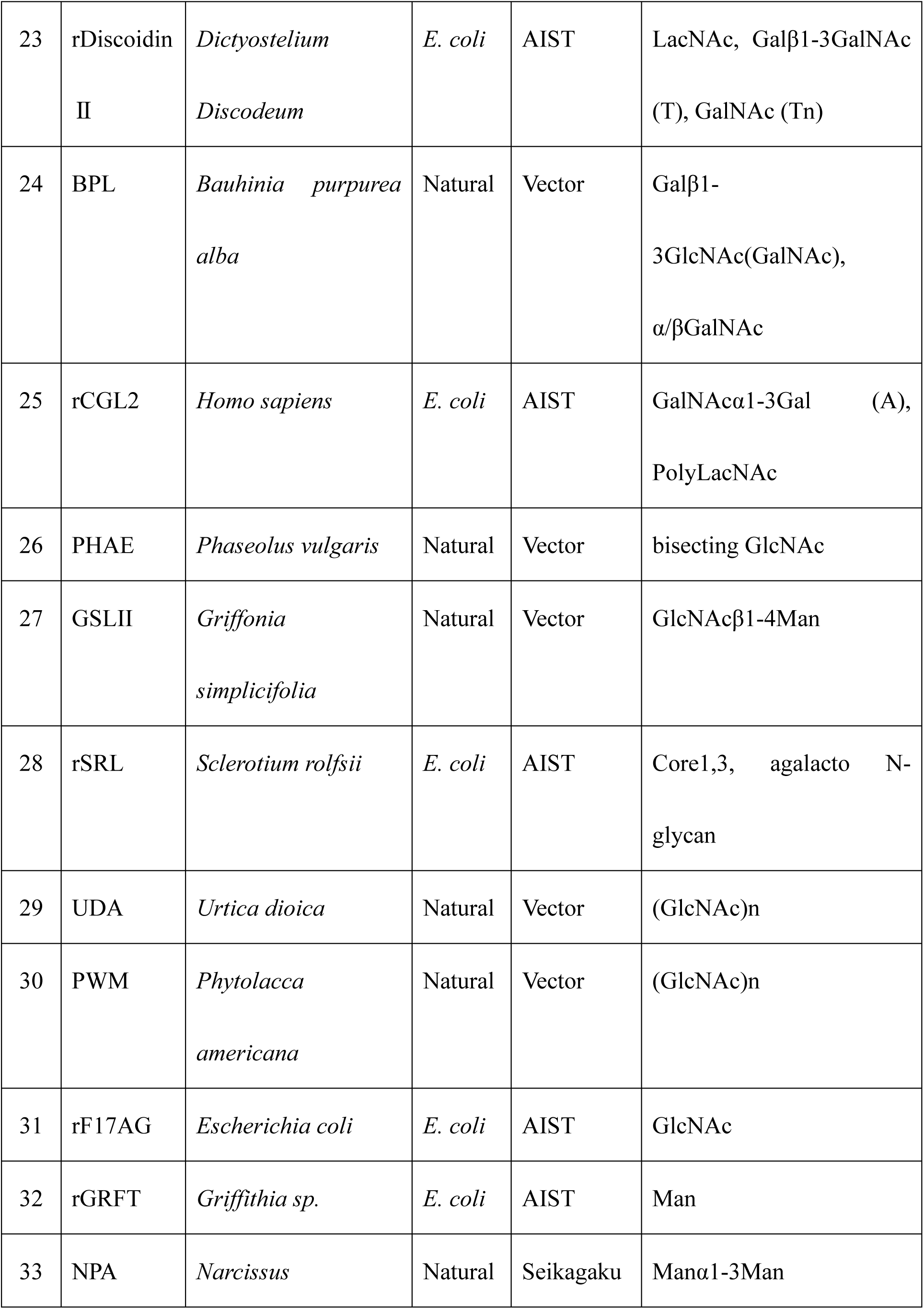
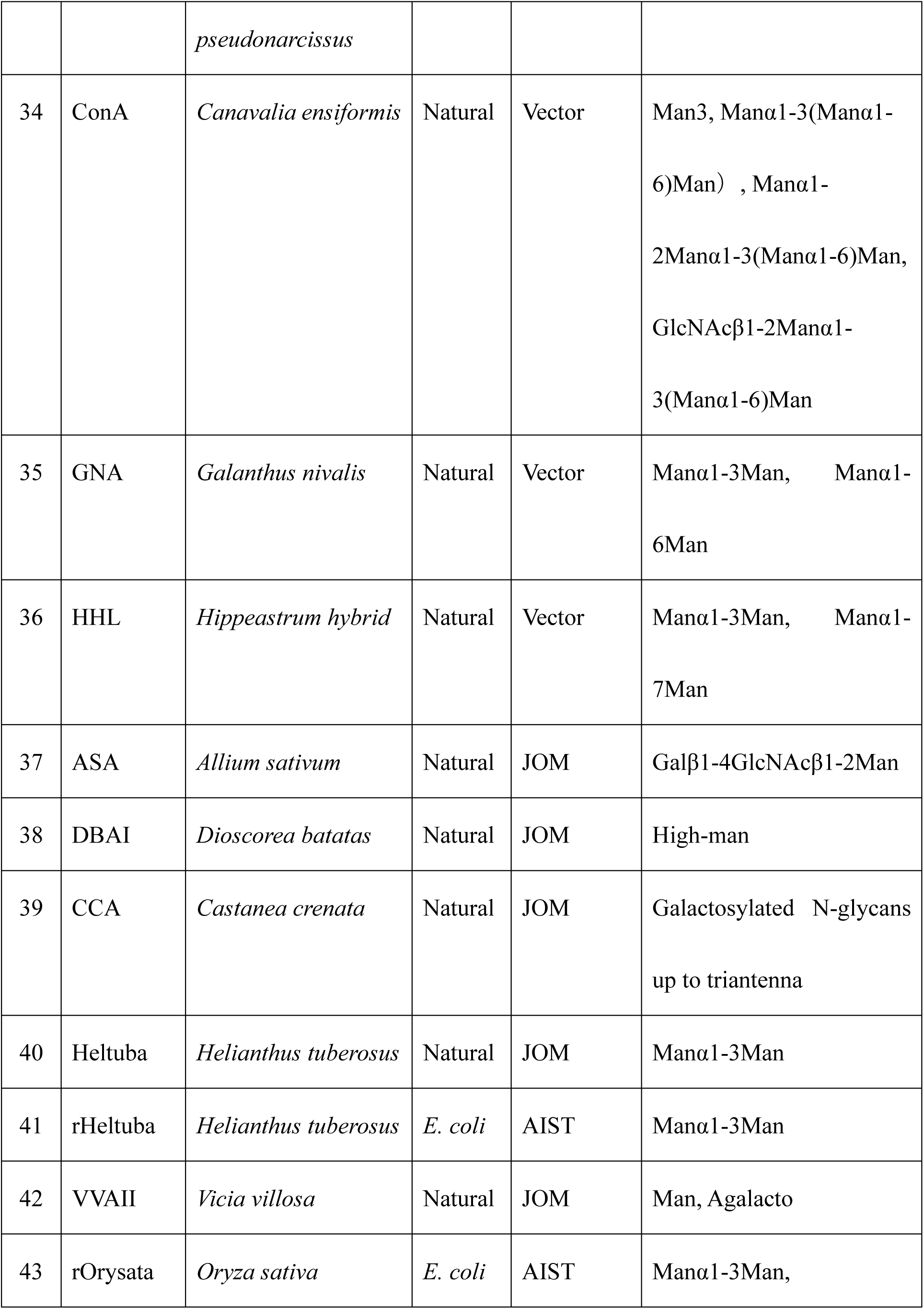
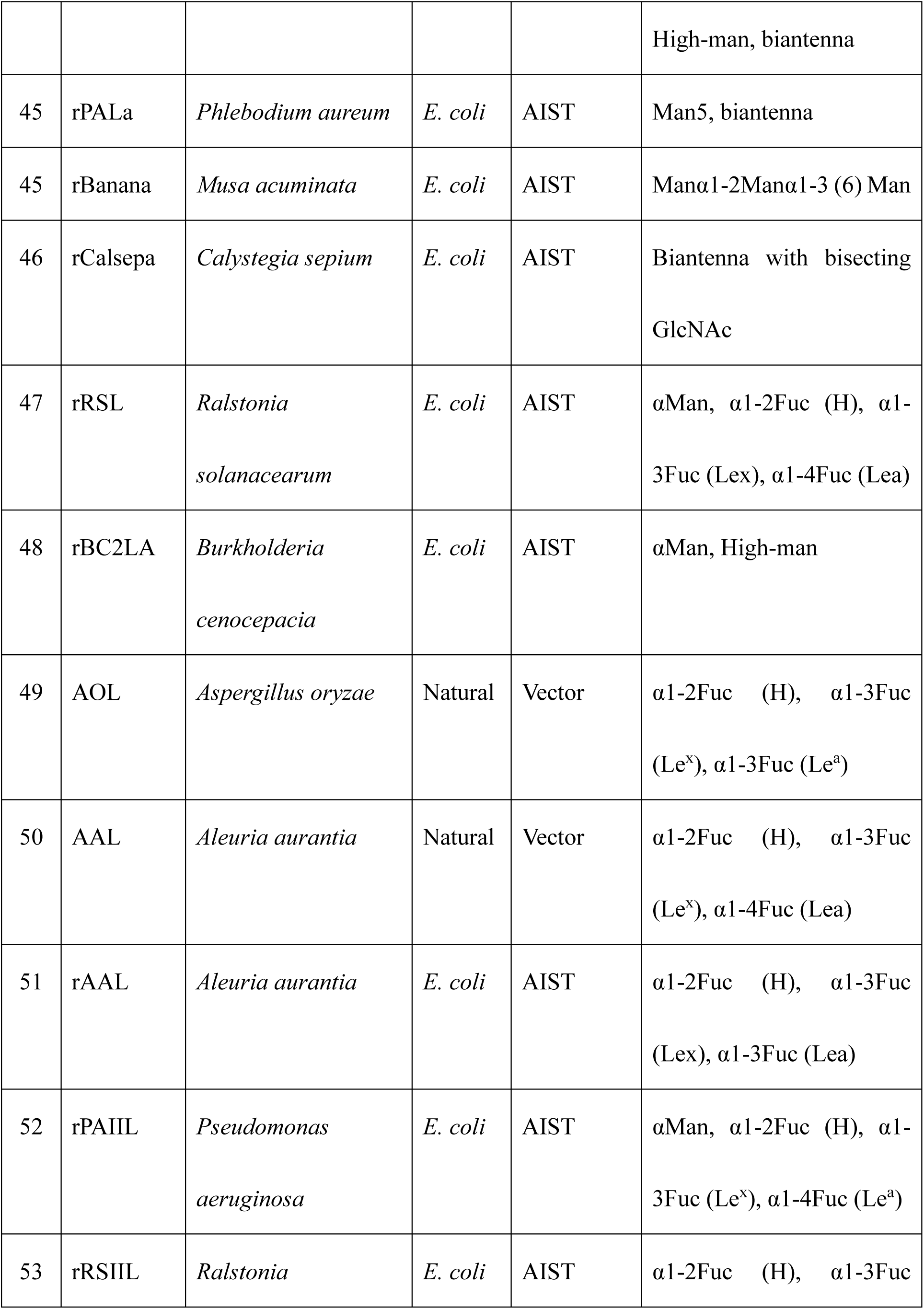
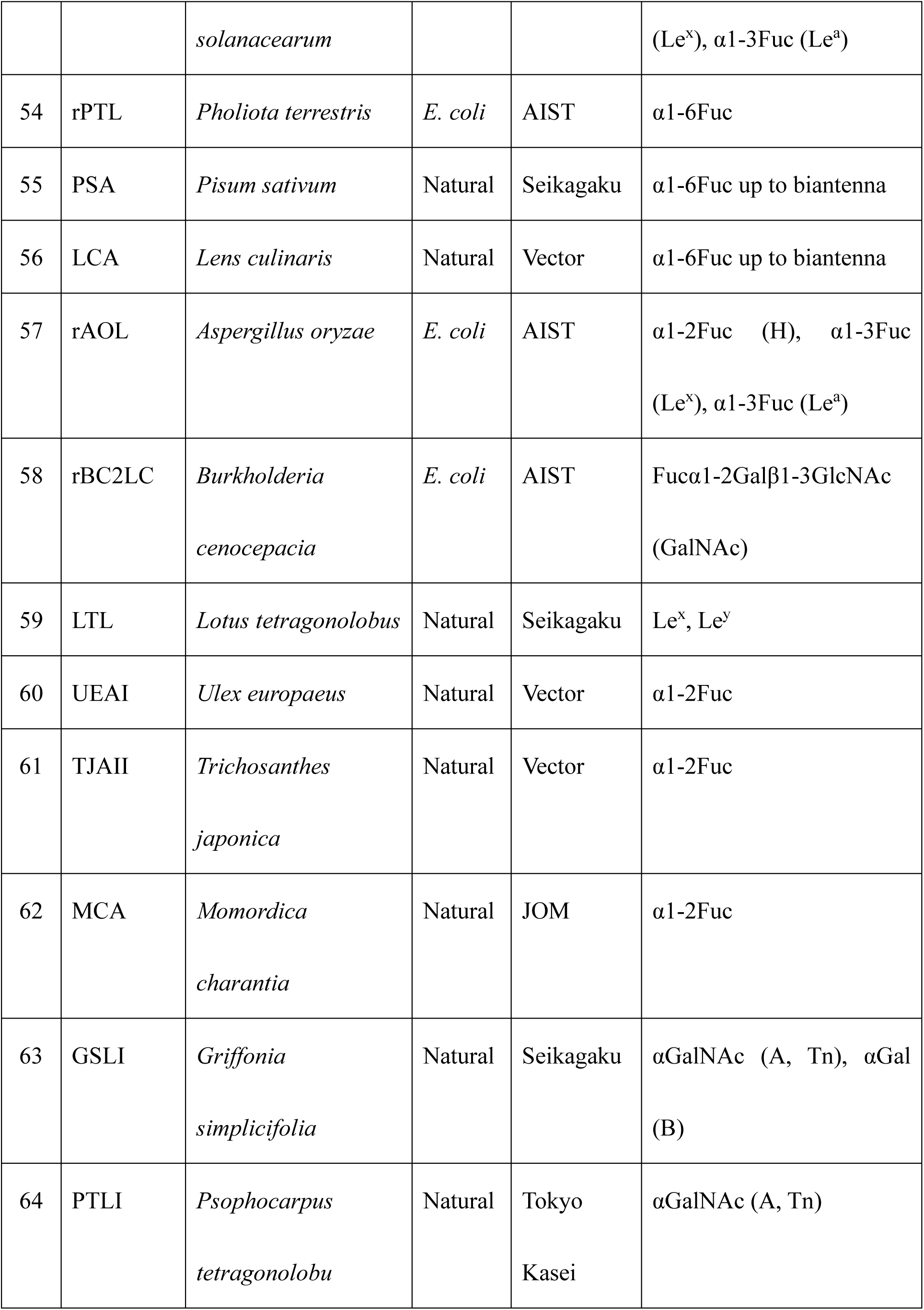
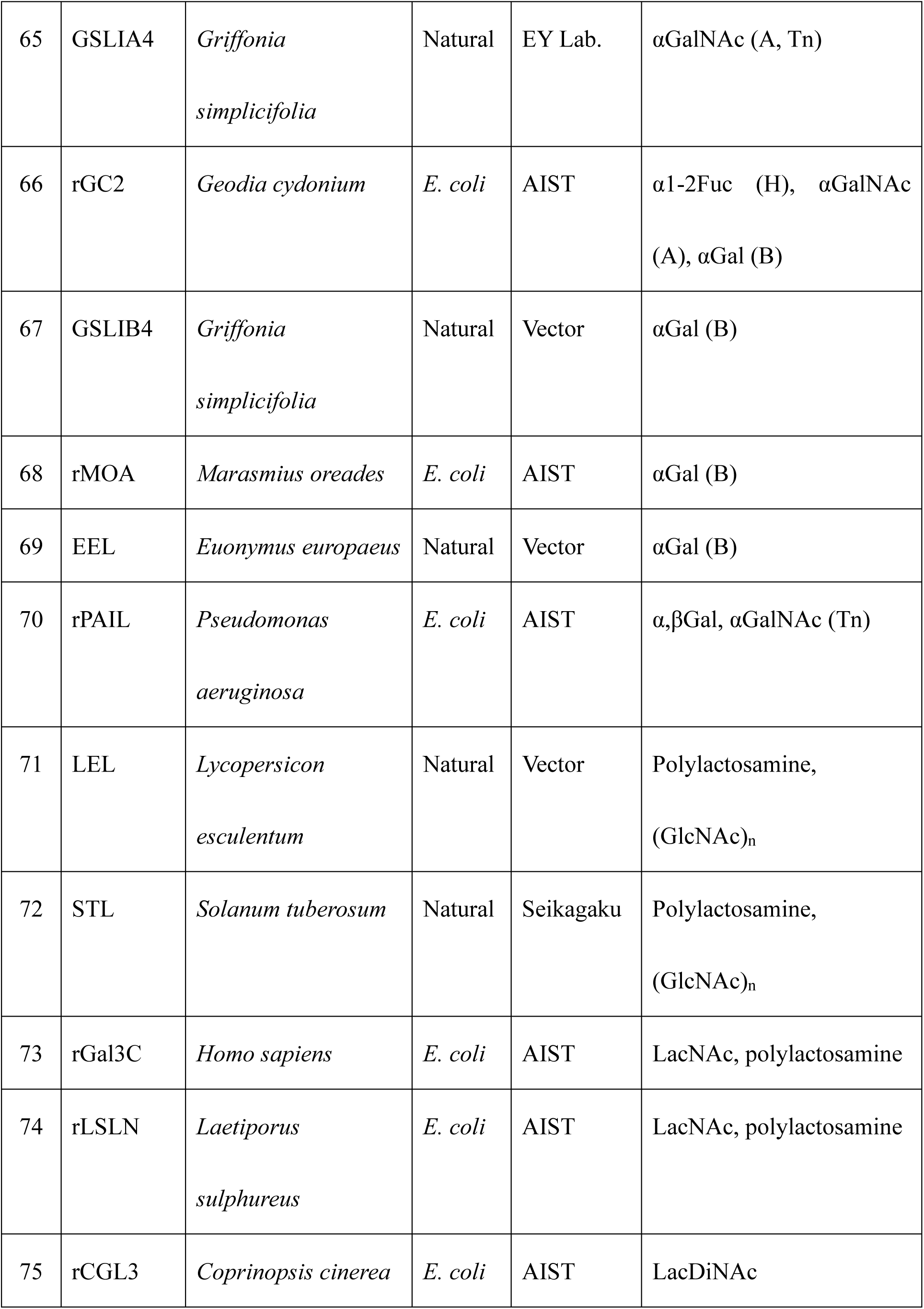
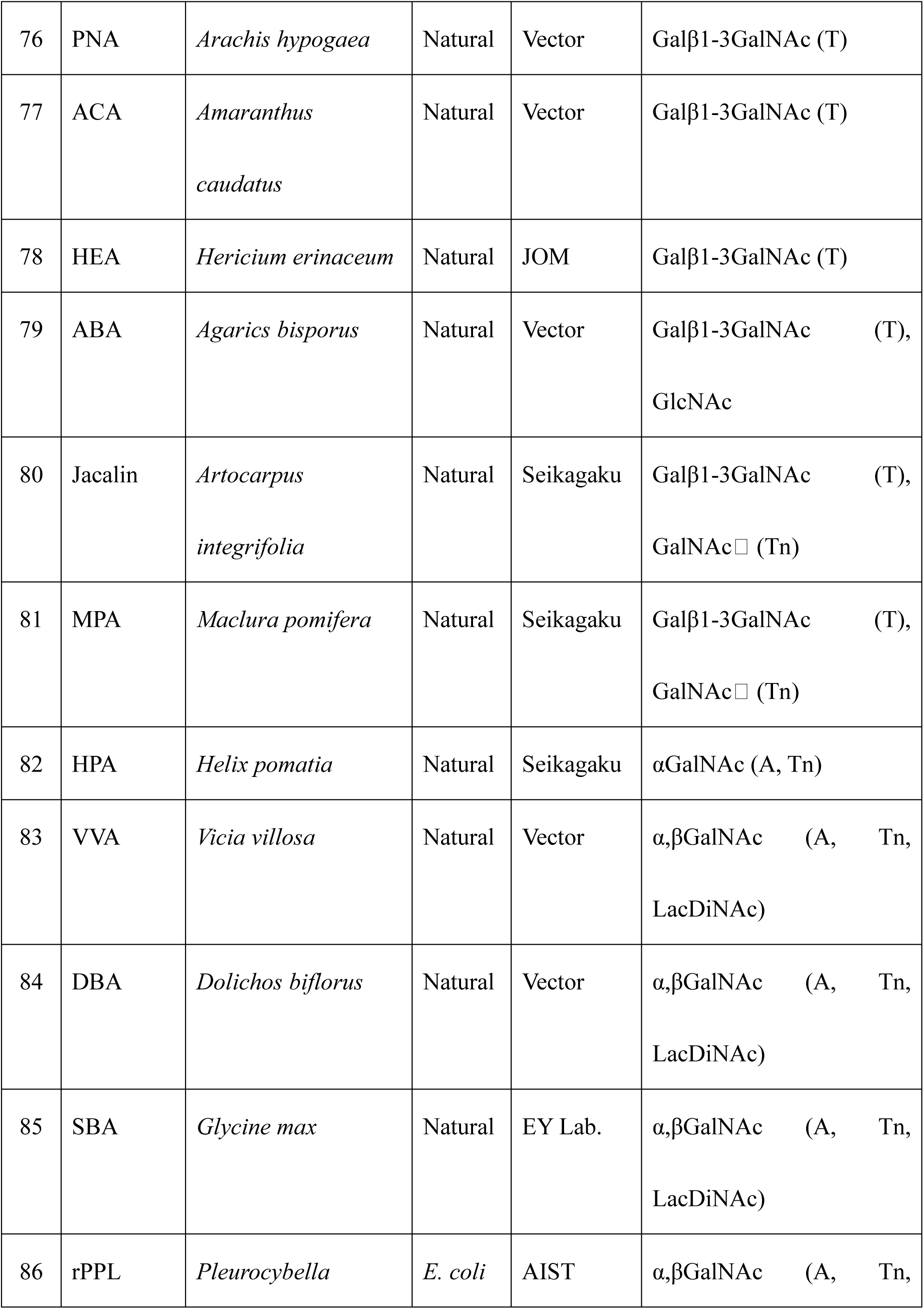
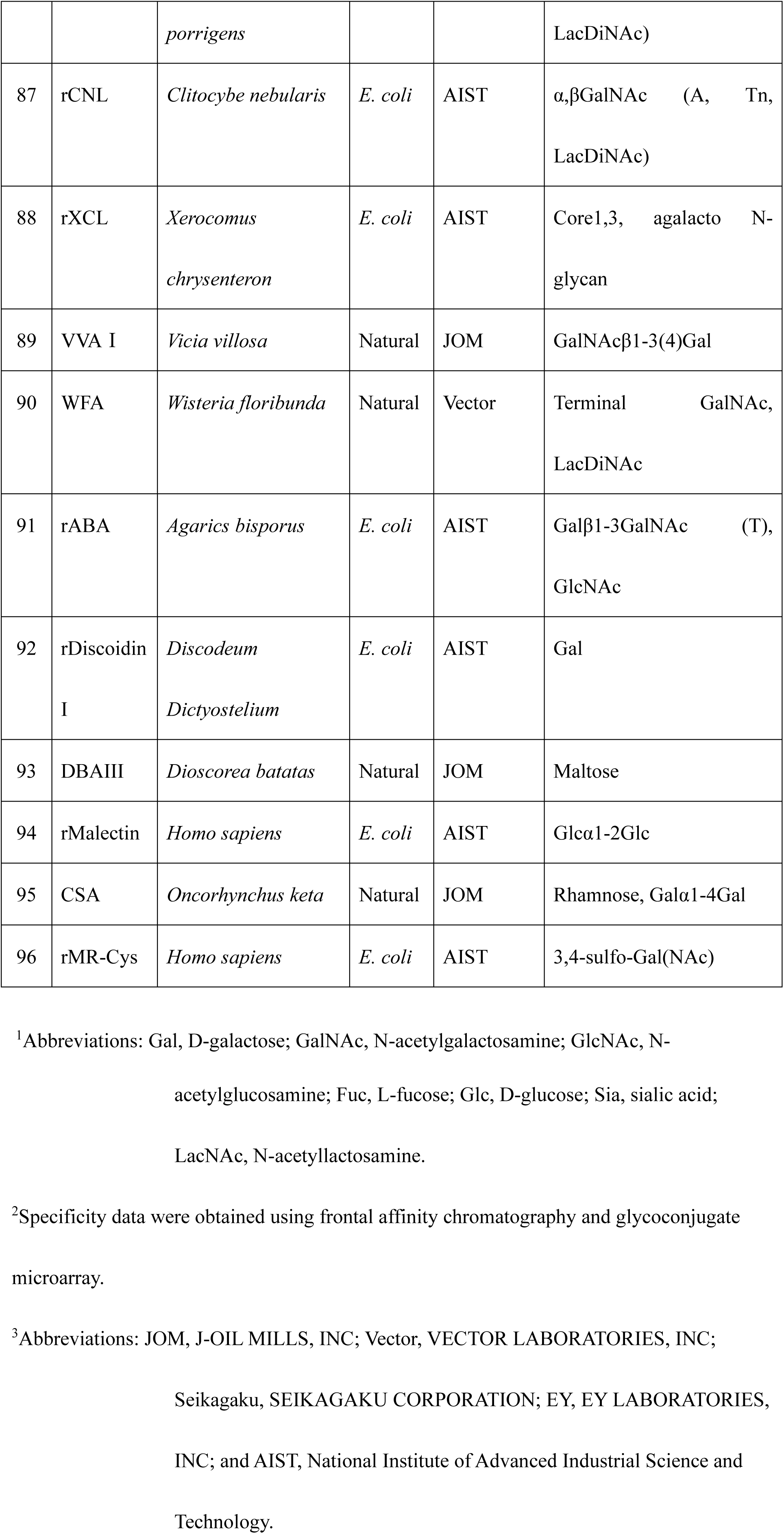
Sugar-binding specificities of the lectins in this study.

## REFERENCES

1. Chapot-Chartier MP, Kulakauskas S. 2014. Cell wall structure and function in lactic acid bacteria. Microb Cell Fact 13.

2. Weintraub A. 2003. Immunology of bacterial polysaccharide antigens. Carbohydr Res. Elsevier BV 10.1016/j.carres.2003.07.008.

3. Mazmanian SK, Kasper DL. 2006. The love-hate relationship between bacterial polysaccharides and the host immune system. Nat Rev Immunol 10.1038/nri1956.

4. Campanero-Rhodes MA, Palma AS, Menéndez M, Solís D. 2020. Microarray Strategies for Exploring Bacterial Surface Glycans and Their Interactions With Glycan-Binding Proteins. Front Microbiol. Frontiers Media S.A. 10.3389/fmicb.2019.02909.

5. Vinogradov E, Sadovskaya I, Grard T, Chapot-Chartier MP. 2016. Structural studies of the rhamnose-rich cell wall polysaccharide of Lactobacillus casei BL23. Carbohydr Res 435:156–161.

6. Zeidan AA, Poulsen VK, Janzen T, Buldo P, Derkx PMF, Øregaard G, Neves AR. 2017. Polysaccharide production by lactic acid bacteria: from genes to industrial applications. FEMS Microbiol Rev. Oxford University Press 10.1093/FEMSRE/FUX017.

7. Shida K, Nanno M, Nagata S. 2011. Flexible cytokine production by macrophages and t cells in response to probiotic bacteria: A possible mechanism by which probiotics exert multifunctional immune regulatory activities. Gut Microbes 2:109–114.

8. Shida K, Nomoto K. 2013. Probiotics as efficient immunopotentiators: Translational role in cancer preventionIndian J Med Res.

9. Cox AJ, Makino H, Cripps AW, West NP. 2019. Recovery of Lactobacillus casei strain Shirota (LcS) from faeces with 14 days of fermented milk supplementation in healthy Australian adults. Asia Pac J Clin Nutr 28:734–739.

10. Takada M, Nishida K, Kataoka-Kato A, Gondo Y, Ishikawa H, Suda K, Kawai M, Hoshi R, Watanabe O, Igarashi T, Kuwano Y, Miyazaki K, Rokutan K. 2016. Probiotic Lactobacillus casei strain Shirota relieves stress-associated symptoms by modulating the gut–brain interaction in human and animal models. Neurogastroenterology and Motility 28:1027–1036.

11. Cook CM, Makino H, Kato K, Blonquist T, Derrig L, Shibata H. 2023. The probiotic Lacticaseibacillus paracasei strain Shirota (LcS) in a fermented milk beverage survives the gastrointestinal tract of generally healthy U.S. Adults. Int J Food Sci Nutr 74:645–653.

12. Otaka M, Kikuchi-Hayakawa H, Ogura J, Ishikawa H, Yomogida Y, Ota M, Hidese S, Ishida I, Aida M, Matsuda K, Kawai M, Yoshida S, Kunugi H. 2021. microorganisms Effect of Lacticaseibacillus paracasei Strain Shirota on Improvement in Depressive Symptoms, and Its Association with Abundance of Actinobacteria in Gut Microbiota 10.3390/microorganisms905.

13. Kikuchi-Hayakawa H, Ishikawa H, Suda K, Gondo Y, Hirasawa G, Nakamura H, Takada M, Kawai M, Matsuda K. 2023. Effects of Lacticaseibacillus paracasei Strain Shirota on Daytime Performance in Healthy Office Workers: A Double-Blind, Randomized, Crossover, Placebo-Controlled Trial. Nutrients 15:5119.

14. Takada M, Nishida K, Gondo Y, Kikuchi-Hayakawa H, Ishikawa H, Suda K, Kawai M, Hoshi R, Kuwano Y, Miyazaki K, Rokutan K. 2017. Beneficial effects of Lactobacillus casei strain Shirota on academic stress-induced sleep disturbance in healthy adults: A double-blind, randomised, placebo-controlled trial. Benef Microbes 8:153–162.

15. Kato-Kataoka A, Nishida K, Takada M, Kawai M, Kikuchi-Hayakawa H, Suda K, Ishikawa H, Gondo Y, Shimizu K, Matsuki T, Kushiro A, Hoshi R, Watanabe O, Igarashi T, Miyazaki K, Kuwano Y, Rokutan K. 2016. Fermented milk containing Lactobacillus casei strain Shirota preserves the diversity of the gut microbiota and relieves abdominal dysfunction in healthy medical students exposed to academic stress. Appl Environ Microbiol 82:3649–3658.

16. Matsumoto K, Takada T, Shimizu K, Kado Y, Kawakami K, Makino I, Yamaoka Y, Hirano K, Nishimura A, Kajimoto O, Nomoto K. 2006. The Effects of a Probiotic Milk Product Containing Lactobacillus casei Strain Shirota on the Defecation Frequency and the Intestinal Microflora of Sub-optimal Health State Volunteers: A Randomized Placebo-controlled Cross-over Study Bioscience Microflora.

17. Takeda K, Okumura K. 2007. Effects of a fermented milk drink containing Lactobacillus casei strain Shirota on the human NK-cell activityJournal of Nutrition. American Institute of Nutrition.

18. Mizukoshi H, Kimura K, Ikemura H, Mori Y, Nagaoka M. 2022. Structural determination of the cell wall polysaccharide LCPS-1 in Lacticaseibacillus paracasei strain Shirota YIT 9029. Carbohydr Res 521.

19. Nagaoka M, Muto M, Nomoto K, Matuzaki T, Watanabe T, Yokokura T. 1990. Structure of Polysaccharide-Peptidoglycan Complex from the Cell Wall of Lactobacillus casei YIT9018J. Biochem.

20. Yasuda E, Serata M, Sako T. 2008. Suppressive effect on activation of macrophages by Lactobacillus casei strain shirota genes determining the synthesis of cell wall-associated polysaccharides, p. 4746–4755. In Applied and Environmental Microbiology.

21. Yuki N, Watanabe K, Mike A, Tagami Y, Tanaka R, Ohwaki M, Morotomi M. 1999. Survival of a probiotic, Lactobacillus casei strain Shirota, in the gastrointestinal tract: Selective isolation from faeces and identification using monoclonal antibodiesInternational Journal of Food Microbiology.

22. Mazé A, Boël G, Zúñiga M, Bourand A, Loux V, Yebra MJ, Monedero V, Correia K, Jacques N, Beaufils S, Poncet S, Joyet P, Milohanic E, Casarégola S, Auffray Y, Pérez-Martínez G, Gibrat JF, Zagorec M, Francke C, Hartke A, Deutscher J. 2010. Complete genome sequence of the probiotic Lactobacillus casei strain BL23. J Bacteriol 192:2647–2648.

23. Dicks LM, Du Plessis EM, Dellaglio F, Lauer E. 1996. Reclassification of Lactobacillus casei subsp. casei ATCC 393 and Lactobacillus rhamnosus ATCC 15820 as Lactobacillus zeae nom. rev., designation of ATCC 334 as the neotype of L. casei subsp. casei, and rejection of the name Lactobacillus paracasei. Int J Syst Bacteriol 46:337–40.

24. Watanabe K, Ishibashi K, Nakashima Y, Sakurai T. 1984. A Phage-resistant Mutant of Lactobacillus casei which Permits Phage Adsorption but Not Genome InjectionJ. gen. Virol.

25. Kiwakil M, Shimizu-Kadotal M. 2002. Development of Genetic Manipulation Systems and the Appli-cation to Genetic Research in Lactobacillus casei Strain Shirota Bioscience Microflora.

26. Yan X, Habbersett RC, Cordek JM, Nolan JP, Yoshida TM, Jett JH, Marrone BL. 2000. Development of a mechanism-based, DNA staining protocol using SYTOX orange nucleic acid stain and DNA fragment sizing flow cytometry. Anal Biochem 286:138–148.

27. Yasuda E, Tateno H, Hirabarashi J, Iino T, Sako T. 2011. Lectin microarray reveals binding profiles of Lactobacillus casei strains in a comprehensive analysis of bacterial cell wall polysaccharides. Appl Environ Microbiol 77:4539–4546.

28. Hirabayashi J, Yamada M, Kuno A, Tateno H. 2013. Lectin microarrays: concept, principle and applications. Chem Soc Rev 42:4443–58.

29. Tateno H, Toyota M, Saito S, Onuma Y, Ito Y, Hiemori K, Fukumura M, Matsushima A, Nakanishi M, Ohnuma K, Akutsu H, Umezawa A, Horimoto K, Hirabayashi J, Asashima M. 2011. Glycome diagnosis of human induced pluripotent stem cells using lectin microarray. Journal of Biological Chemistry 286:20345–20353.

30. W Zhang WJPABCHAPRPRPPPT-BAAJEJVD. 2000. Isolation and characterization of a jacalin-related mannose-binding lectin from salt-stressed rice (Oryza sativa) plants. Planta 210:970–978.

31. Nagae M, Mishra SK, Hanashima S, Tateno H, Yamaguchi Y. 2017. Distinct roles for each N-glycan branch interacting with mannose-binding type Jacalin-related lectins Orysata and Calsepa. Glycobiology 27:1120–1133.

32. Barondes SH, Cooper DN, Haywood-Reid PL. 1983. Discoidin land Discoidin II Are Localized in Developing Dictyostelium discoideum Differently TH£ JOURNAL OF CELL BIOLOGY.

33. Mistou MY, Sutcliffe IC, Van Sorge NM. 2016. Bacterial glycobiology: Rhamnose-containing cell wall polysaccharides in gram-positive bacteria. FEMS Microbiol Rev. Oxford University Press 10.1093/femsre/fuw006.

34. Graninger M, Kneidinger B, Bruno K, Scheberl A, Messner P. 2002. Homologs of the Rml enzymes from Salmonella enterica are responsible for dTDP-β-L-rhamnose biosynthesis in the gram-positive thermophile Aneurinibacillus thermoaerophilus DSM 10155. Appl Environ Microbiol 68:3708–3715.

35. Singh SS, Devi SK, Ng TB. 2014. Banana lectin: A brief review. Molecules. MDPI AG 10.3390/molecules191118817.

36. Mo H, Winter HC, Van Damme EJM, Peumans WJ, Misaki A, Goldstein IJ. 2001. Carbohydrate binding properties of banana (Musa acuminata) lectin: I. Novel recognition of internal α1,3-linked glucosyl residues. Eur J Biochem 268:2609–2615.

37. Naito Y, Minamihara T, Ando A, Marutani T, Oguri S, Nagata Y. 2001. Domain construction of cherry-tomato lectin: relation to newly found 42-kDa protein. Biosci Biotechnol Biochem 65:86–93.

38. Parasuraman P, Murugan V, Selvin JFA, Gromiha MM, Fukui K, Veluraja K. 2014. Insights into the binding specificity of wild type and mutated wheat germ agglutinin towards Neu5Acα(2-3)Gal: a study by in silico mutations and molecular dynamics simulations. J Mol Recognit 27:482–92.

39. Itakura Y, Nakamura-Tsuruta S, Kominami J, Tateno H, Hirabayashi J. 2017. Sugar-binding profiles of chitin-binding lectins from the hevein family: A comprehensive study. Int J Mol Sci 18.

40. Schallus T, Jaeckh C, Fehér K, Palma AS, Liu Y, Simpson JC, Mackeen M, Stier G, Gibson TJ, Feizi T, Pieler T, Muhle-Goll C. 2008. Malectin: a novel carbohydrate-binding protein of the endoplasmic reticulum and a candidate player in the early steps of protein N-glycosylation. Mol Biol Cell 19:3404–14.

41. Shirai T, Watanabe Y, Lee M sub, Ogawa T, Muramoto K. 2009. Structure of Rhamnose-binding Lectin CSL3: Unique Pseudo-tetrameric Architecture of a Pattern Recognition Protein. J Mol Biol 391:390–403.

42. Shimizu-Kadota M, Kiwaki M, Sawaki S, Shirasawa Y, Shibahara-Sone H, Sako T. 2000. Insertion of bacteriophage wFSW into the chromosome of Lactobacillus casei strain Shirota (S-1): Characterization of the attachment sites and the integrase gene Gene.

